# Dopamine neurons encode a multidimensional probabilistic map of future reward

**DOI:** 10.1101/2023.11.12.566727

**Authors:** Margarida Sousa, Pawel Bujalski, Bruno F. Cruz, Kenway Louie, Daniel McNamee, Joseph J. Paton

## Abstract

Learning to predict rewards is a fundamental driver of adaptive behavior. Midbrain dopamine neurons (DANs) play a key role in such learning by signaling reward prediction errors (RPEs) that teach recipient circuits about expected rewards given current circumstances and actions. However, the algorithm that DANs are thought to provide a substrate for, temporal difference (TD) reinforcement learning (RL), learns the mean of temporally discounted expected future rewards, discarding useful information concerning experienced distributions of reward amounts and delays. Here we present time-magnitude RL (TMRL), a multidimensional variant of distributional reinforcement learning that learns the joint distribution of future rewards over time and magnitude using an efficient code that adapts to environmental statistics. In addition, we discovered signatures of TMRL-like computations in the activity of optogenetically identified DANs in mice during a classical conditioning task. Specifically, we found significant diversity in both temporal discounting and tuning for the magnitude of rewards across DANs, features that allow the computation of a two dimensional, probabilistic map of future rewards from just 450ms of neural activity recorded from a population of DANs in response to a reward-predictive cue. In addition, reward time predictions derived from this population code correlated with the timing of anticipatory behavior, suggesting the information is used to guide decisions regarding when to act. Finally, by simulating behavior in a foraging environment, we highlight benefits of access to a joint probability distribution of reward over time and magnitude in the face of dynamic reward landscapes and internal physiological need states. These findings demonstrate surprisingly rich probabilistic reward information that is learned and communicated to DANs, and suggest a simple, local-in-time extension of TD learning algorithms that explains how such information may be acquired and computed.

## Introduction

The field of reinforcement learning (RL) provides a normative theoretical framework for adaptive animal behavior^2^. A core tenet of RL is that behaviors producing maximal expected future reward are the target of learning through interaction with the environment. Relatedly, associative learning, including learning to associate states and actions with future reward as is required by RL, can be viewed through the lens of statistical inference^3^. Whether a given observation or action should be associated with future reward depends on the degree to which its taking place reduces uncertainty about when rewards will occur. Such an account for associative learning would seem to require the brain to operate on distributions of events in time^4^. Furthermore, predicting when, and not just whether, behaviorally relevant events such as rewards will occur is often critical for survival. For example, crossing a desert to reach an oasis is only advisable if you won’t perish before you get there. However, the RL algorithms that have driven startling progress in the neuroscience of learned behavioral control and in artificial intelligence alike, do not generally learn value representations that encode distributions of rewards over time.

Midbrain dopamine neurons have figured prominently in theories of how RL-like functions may be performed within neural circuits. Specifically, the phasic activity of midbrain DANs is thought to encode temporal difference (TD) reward prediction errors (RPEs), which serve as teaching signals that are used to update the value of states or actions so as to inform appropriate programs, or policies, for behavioral control^1,5^. However, standard TD formulations learn value functions that encode expectations of, temporally discounted, future reward: delayed rewards are weighted less relative to immediate ones. This produces ambiguity in value representations regarding when future rewards are expected to arrive. To illustrate this ambiguity one may observe the same value corresponding to either a large magnitude but delayed reward, or a smaller magnitude but imminent reward (Figure 1A). In addition, because simple TD learning algorithms learn the average of temporally discounted future reward, they do not learn about the distribution of reward magnitudes. Recently, TD algorithms have been elaborated to include a set of value learning channels that differ in their sensitivity to positive and negative RPEs, leading to value estimates that converge to distinct statistics of the expected cumulative reward distribution, an innovation termed distributional RL^6^ (Figure 1B). Such innovations have been shown to improve performance of deep RL agents on benchmark tasks due to improved statistical robustness, and evidence of distributional RL-like computations has been reported in midbrain DANs of both mice and primates^7^,^8^,^9^, however the direct functional relevance of such distributional mechanisms and representations to behavior is unknown. In the engineering setting, deep RL agents vary in whether and how they make use of knowledge about the distribution over future rewards when selecting actions^10–12^, and decoding of reward distributions from DAN activity has only been demonstrated at the time of reward delivery ^7^. In principle, knowing in advance, at the start of an episode, about the range and likelihood of rewards available and when they are likely to occur could be highly useful for planning and flexible behavior, particularly in the face of non-stationary in either the environment or internal state (e.g. hunger) of the animal ^13^.

**Figure 1:**
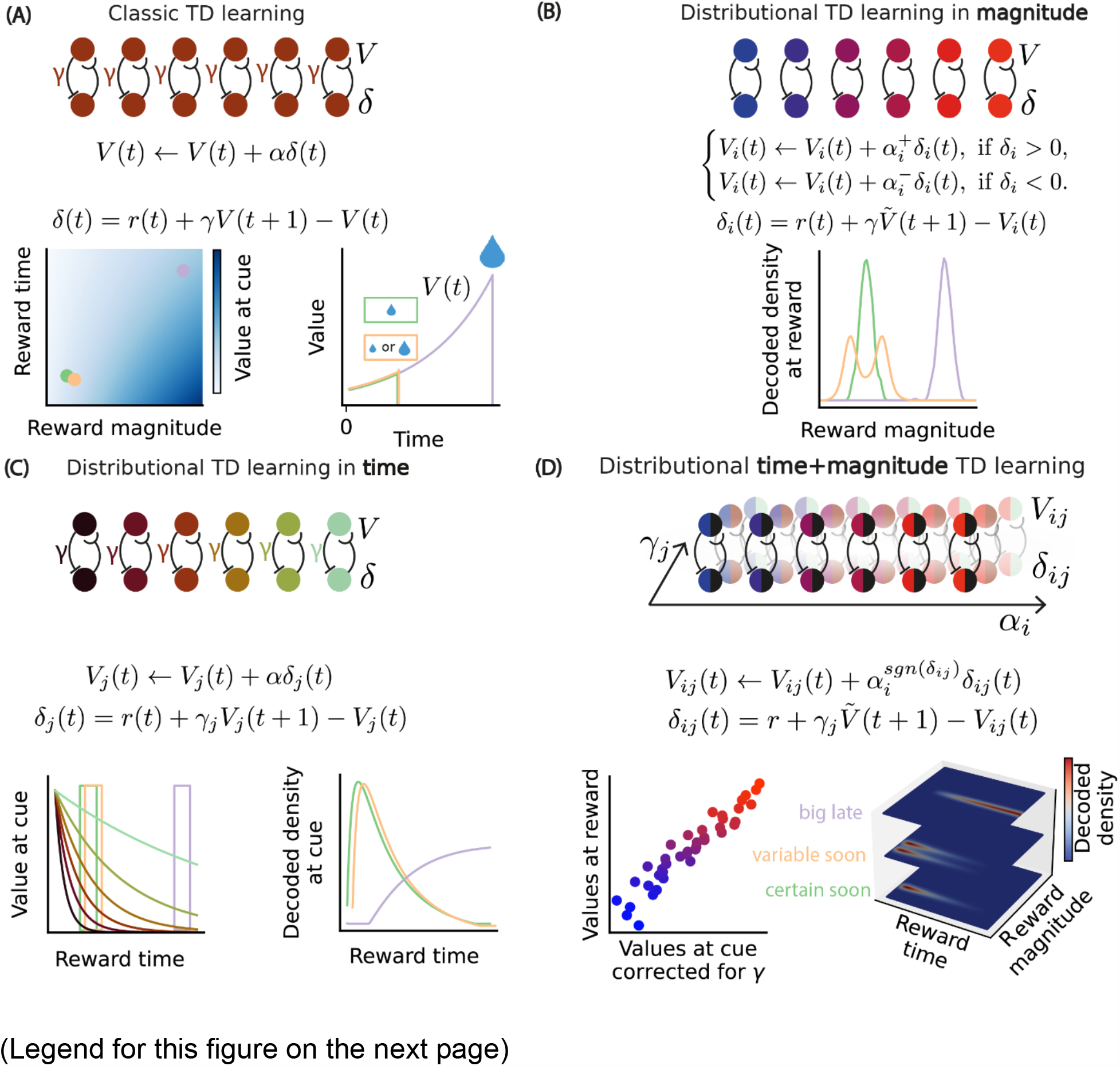
Diversity in temporal discounting and relative scaling for positive and negative RPEs facilitates the construction of a distributional map of future reward in time and magnitude. ***(A)*** *The green cue predicts a certain reward amount after a short delay, the orange a variable amount after a short delay and the purple a big amount after a long delay. In classic temporal-difference (TD) learning all units are computing the RPE relative to the same value, and hence at the beginning of the episode the population cannot distinguish between the three cues*. ***Bottom left:*** *The value at the cue for different reward magnitudes and times. Different combinations of reward magnitude and time map to one value*. ***Bottom right:*** *The temporally discounted value for the three different cues as a function of time since cue*. ***(B)*** *In value distributional TD learning, units asymmetrically scale positive and negative RPEs and hence learn a diverse set of values. We use the learnt values (expectiles) and the asymmetries to decode at the time of reward the distribution over reward magnitudes represented on the bottom*. ***(C)*** *A population with diversity in temporal discount factors allows for decoding the distribution over future rewards at the cue*. ***Bottom left:*** *multiple temporally discounted values, color coded by temporal discount factor. The orange, green and purple blocks represent the population responses at the three different cues*. ***Bottom right:*** *using the population responses at the cue, and having knowledge of the temporal discount factor of each unit, we decoded the future reward distribution over reward time for the three different cues*. ***(D)*** *Considering a population with diverse temporal discount factors and values allows for decoding the map of future reward in time and amount at the cue*. ***Bottom left:*** *Simulated values encoded at reward time as a function of values encoded at the cue corrected for the diversity in temporal discount factor. Points are color coded by value*. ***Bottom right:*** *We use the asymmetries for positive and negative RPEs, the temporal discount factors and the responses at the cue to decode the* probability of reward over time and magnitude.

Here, we develop a computational model of efficient multidimensional distributional RL that learns to predict distributions of rewards over both time and magnitude or distributional time-magnitude RL (TMRL). We then test predictions of the model *in vivo* by recording from optogenetically identified midbrain DANs in mice during behavior. Specifically, in addition to previously demonstrated variability in the degree to which individual neurons respond to positive and negative RPEs ^7^ the model predicts variability in the degree to which individual neurons discount future rewards. We discovered evidence of both types of heterogeneity in DANs. This enabled decoding of future reward times that correlated with the variability in the temporal evolution of behavior, suggesting that decoded estimates correspond with animals’ temporal expectations. Furthermore, we show that taking into account the variability in both sensitivity to reward magnitude and delay allows decoding of a two-dimensional distribution, or map, of future reward amount over time from a set of DAN’ responses to a predictive cue, at the start of an episode. Strikingly, the tuning of individual dopamine neurons for reward time and magnitude was dynamic, adapting to changes in reward statistics, in accordance with the principles of an efficient code that is optimized for encoding information about these two important dimensions of reward. We propose that these data reflect a mechanism by which the brain may use a local-in-time algorithm akin to TD learning to build information-maximising, predictive, and probabilistic models of the environment for use in behavioral control.

## Results

### Learning and encoding a two-dimensional probabilistic map of future reward

We begin by defining an adaptive distributional code for reward time and magnitude (TMRL). This code takes inspiration from several threads of research on temporal discounting^14^, temporal coding in general^15^, distributional value codes, and more recent work that extends distributional value codes to the time domain^7,16,17^.

Classical TD learning produces a global value function that encodes the average of expected future rewards. Temporal discounting of future rewards arises at the computation of the TD RPE *δ(t)* - the temporal difference between the value at the current timestep *V*(*t*) and the discounted value, parameterized by a discount factor *γ*, at the subsequent step *V*(*t + 1*), plus any incoming reward *r*(*t*),

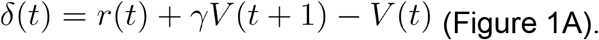

Though expected reward delays are indeed reflected in the value function because of temporal discounting, delay and magnitude information are compressed into a single scalar value, producing ambiguity between these two dimensions in the value code (Figure 1A bottom). Instead of learning a single value function, TD algorithms have recently been elaborated to learn a set of value functions *V*_*i*_ (*t*) that systematically differ in their sensitivity to positive and negative RPEs (*α* ^+^and *α*^−^, Figure 1B),

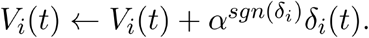

This causes each value function to converge to a different statistic of the observed distribution of reward magnitudes. Viewed collectively, this set of value functions encodes not just the average magnitude of expected future reward, but the distribution over their magnitudes (distributional TD learning in magnitude, Figure 1B). A central insight of our model is that this approach may be generalized to both learn and encode information about the distribution of reward times by assuming that a set of value functions *V*_*j*_ (*t*) are learned that differ in their sensitivity to reward delays, parameterized as the standard temporal discount factor *γ*_*j*_ within the computation of a TD RPE (distributional TD learning in time, Figure 1C),

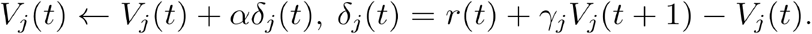

Multiple timescales for discounting across a set of parallel learning channels alone resolves ambiguity between reward delays and the average reward magnitude that is present in a system with a single temporal discount factor (Figure 1C). However, when distributional learning in time is combined with distributional learning of reward magnitude,

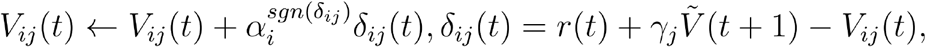

the resulting system learns to encode a probabilistic map of future rewards over both dimensions (Figure 1D). In such a two-dimensional distributional reward coding system, correcting for tuning diversity across one dimension should reveal the remaining tuning diversity for the other that might otherwise be obscured (Figure 1D bottom left).

Critically, because it specifies how temporal and magnitude parameters are learned, the TMRL model adapts to the rewards it experiences. The brain contains a finite number of neurons, and thus faces constraints in its information encoding capacity^18^. In the face of such constraints, efficient coding theory prescribes that the tuning properties of neurons adapt to the statistics of the variable they aim to represent in a manner that maximises overall information content of encoded signals^19,20^. In the current context, this predicts that the discount factors and value parameters should be adapted so as to maximise the encoding of reward time-magnitude information with respect to the current environment^21^. We return to this aspect of the TMRL model in more detail below.

### Temporal discount rates vary among dopamine neurons and carry information about the distribution of future reward times

Does the brain use an algorithm similar to TMRL to learn about distributions of rewards along multiple dimensions? We tested for this by recording from midbrain DANs of mice (Ext. Data Fig. 1 and 2) during a simple behavioral task, trace odor conditioning, designed to induce predictions of reward at different delays and magnitudes. Four odor cues (conditioned stimuli, CSs) predicted the same reward amount but with a distinct delay (0,1.5, 3 or 6 seconds^a^, respectively, Figure 2A). A fifth CS predicted, at a delay of 3s, a reward amount sampled from a bimodal probability distribution (Figure 2B). Importantly the mean of the probability distribution of rewards associated with this fifth CS was equal to the fixed reward amount delivered following presentation of the other CSs. We focus our analyses on data from 43 optogenetically identified DANs collected from 6 trained mice. By considering the response of each dopamine neuron to CSs with different delays to reward, we estimated how single neurons discounted rewards over time, parametrizing this function with a temporal discount factor (*γ*), and a gain parameter (Figure 2C). In addition, by examining the responses to different reward amounts, we estimated the value expected by each neuron (reversal point) and the slopes for negative (*α*^−^, represented in blue) and positive (*α* ^+^, represented in red) RPEs ^7^, Figure 2D).

**Figure 2:**
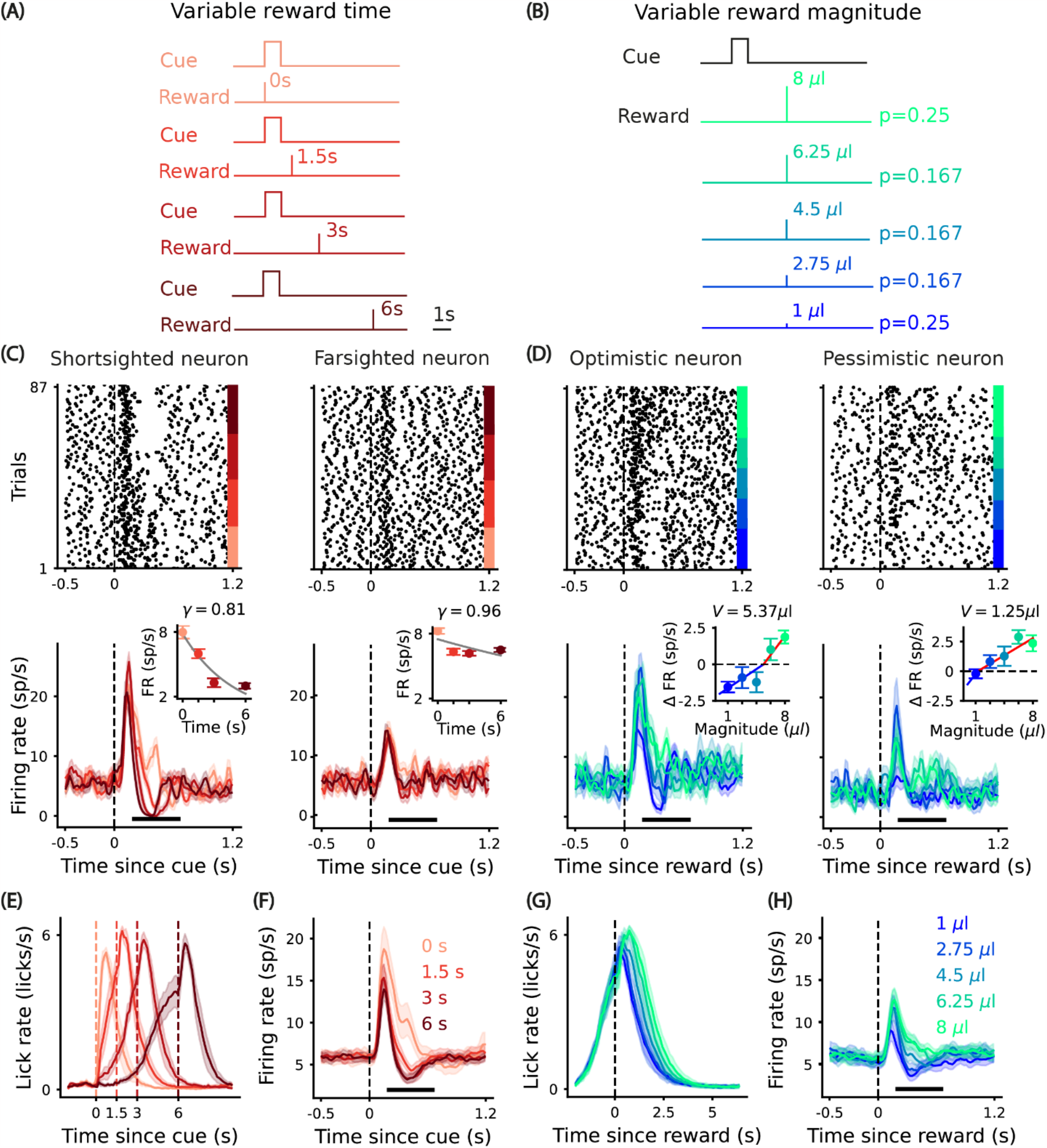
*Dopamine neurons are modulated by reward magnitude and time*. ***(A)*** *Variable time CSs, trial types 1-4, odor cues are sampled to produce a uniform distribution of reward times over trials, reward magnitude = 4*.*5 μl*. ***(B)*** *Variable magnitude CSs, trial type 5. 3s after CS onset, a reward amount sampled from a bimodal distribution is delivered*. ***(C)*** *Raster and* mean peristimulus histogram (*PSTH) aligned to odor onset for different reward times for two example neurons. The shaded area depicts the standard error of the mean. Black line: window used to compute responses (200-650ms). The inset depicts responses to the different delays, gray line: fitted discount function*. ***(D)*** *Raster and PSTH aligned to reward delivery for different reward magnitudes. The inset for the PSTH depicts the baseline subtracted responses for the five different reward amounts, blue line: fitted line for the negative responses, red line: fitted line for the positive responses*. ***(E)*** *Mean lick rate for all sessions and all animals (n=6) aligned to odor onset for different reward times, the shaded area depicts the standard error of the mean*. ***(F)*** *Mean population PSTH aligned to odor onset, the shaded area depicts the standard error of the mean*. ***(G)*** *Mean licking rate for all sessions and all animals (n=6) aligned to reward delivery for different reward magnitudes, the shaded area depicts the standard error of the mean*. ***(H)*** *Mean population PSTH aligned to reward delivery, the shaded area depicts the standard error of the mean*.

We tested for the diversity in temporal discounts in optogenetically identified dopamine neurons and in putative dopamine neurons (see Ext. Data Fig. 3 and 4). We identified significant diversity in temporal discounts (Figure 3A-C). This diversity did not reflect noise in the estimation of *γ*, as estimates were highly correlated across random partitioning of trials (Figure 3B). Furthermore, neurons exhibited significantly different temporal discounts (Figure 3C). We note that five cells possessed estimated discount factors that were greater than, and with 99% confidence intervals that were non-inclusive of, one. This may reflect limitations due to a limited number of reward delays, since three of these neurons were recorded for the mice only subject to three (instead of four) reward delays; by considering an underestimate of the responses to the missing delay, we obtain temporal discount factors that do not significantly differ from one (see Ext. Data Fig. 5). For completeness we include these neurons in all analyses.

**Figure 3:**
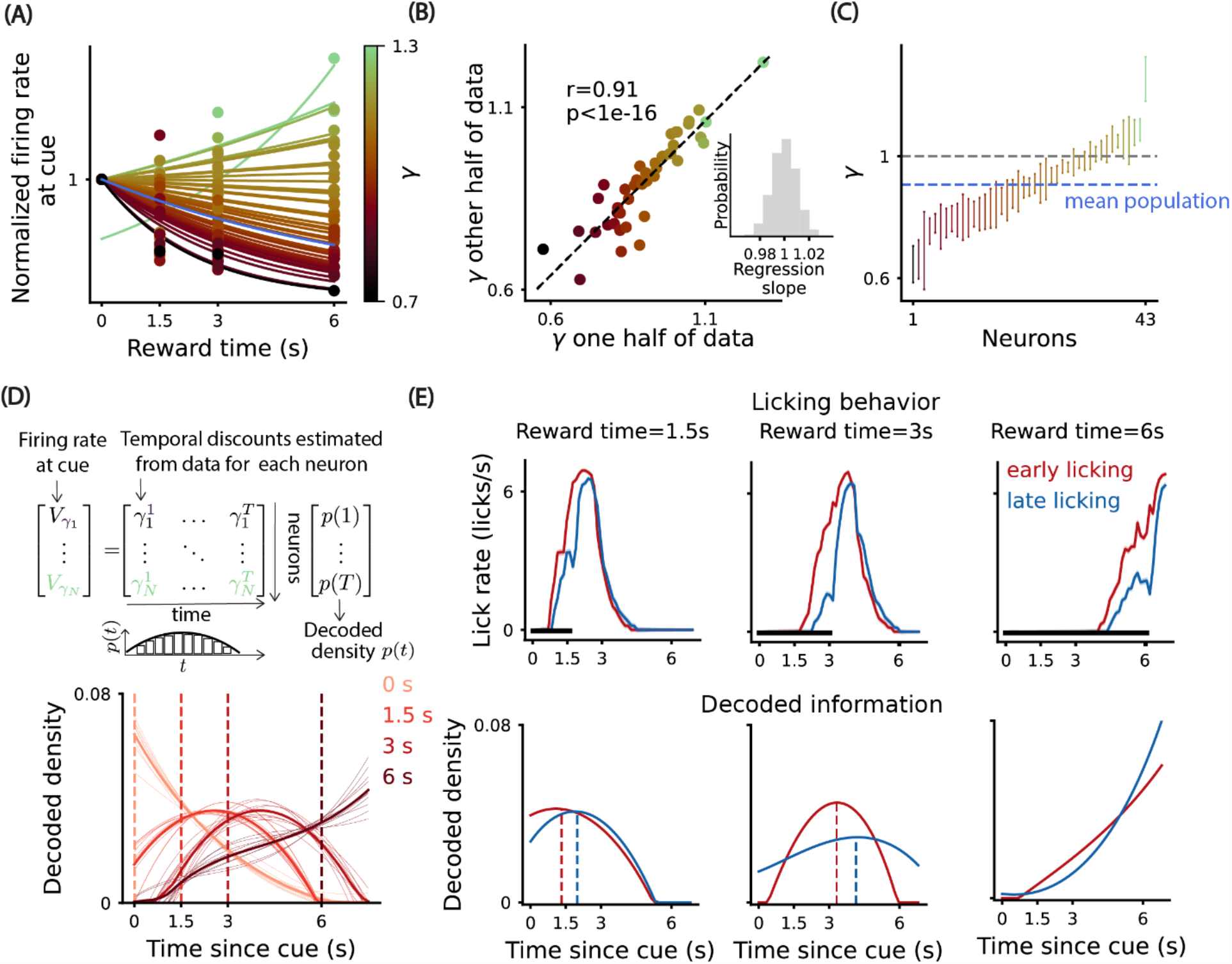
*Dopamine neurons discount future reward heterogeneously, reflecting information about the timing of future rewards that correlates with licking behavior*. ***(A)*** *The dots are single neuron responses to CSs predicting different reward times, normalized by the responses at delay=0s and the lines are the fitted temporal discount functions, color code: temporal discount factor. The population mean temporal discount function is plotted in blue*. ***(B)*** *Temporal discounts estimated using two random disjoint partitions of the total number of trials. This procedure was repeated 10 000 times, and the mean correlation coefficient and the geometric mean of the p-values (two-tailed test) is depicted. In the inset, the histogram of the regression slopes for each run is depicted*. ***(C)*** *Cross validation of temporal discount factors of single neurons using 50% of the total number of trials per delay. The error bars: 99% confidence interval. Dashed blue line: population mean temporal discount factor, dashed gray line: temporal discount factor equal to one*. One-way ANOVA for the difference in population temporal discounts F(42,252)=1498.02, p-value=1.96e-284. ***(D) Top:*** *We use the responses at the cue (normalized by the gain) and the estimated temporal discount for each neuron to decode the distribution of future reward by inverting a linear regression*. ***Middle:*** *A representation of a possible decoded density over a discretized range of times*. ***Bottom:*** *decoded density using the dopamine population responses aligned to odor cue onsets predicting rewards after 0s, 1*.*5, 3s and 6s respectively. The light lines represent decoded densities using the responses of 70% of randomly selected trials and the dark lines represent the mean decoded density*. ***(E) Top:*** *Mean licking rate for all mice (n=6), for the trials in which the mice started licking earlier (red) or later (blue). The shaded area depicts the standard error of the mean and the horizontal black line the window used to compute the licking slopes*. ***Bottom:*** *Decoded density using the trials for which the mice started licking earlier (red) or later (blue). The dashed lines depict the maximum of the decoded reward time*.

**Figure 4:**
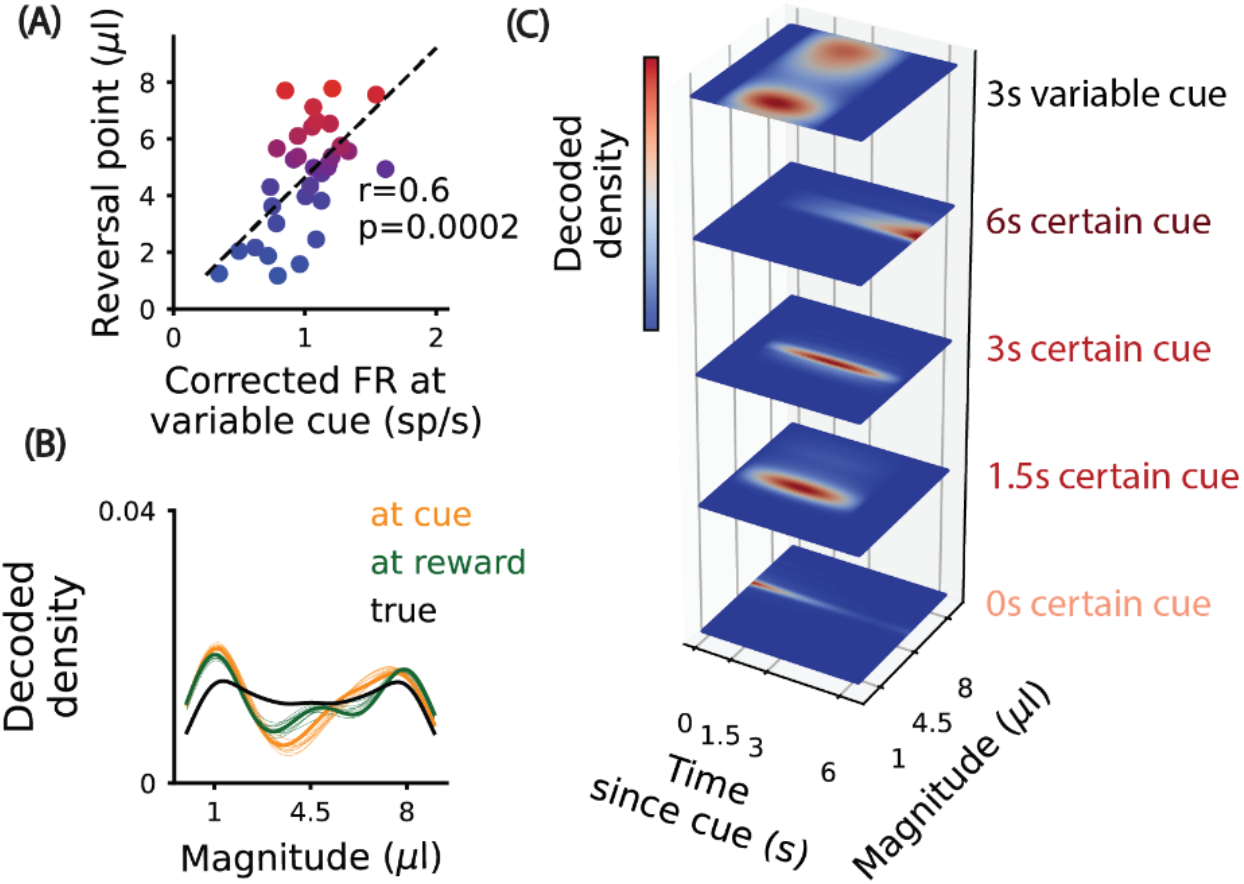
Dopamine neurons reflect information about the distribution of future rewards at cue presentation. **(A)** *Reversal points of individual neurons (calculated from the response to rewards of differing amount) as a function of the response to the cue associated with variable reward, corrected for the estimated discount function for each neuron (calculated from the response to cues associated with rewards of differing delay), color code: reversal point*. Pearson correlation coefficient r=0.6, two-tailed test p-value=0.0002, 95% CI=(0.33,0.79). ***(B)*** *The actual smoothed distribution over reward amounts predicted by the variable cue (black) and the density over reward amounts decoded from the DAN population response to reward (green) and the cue predicting variable reward amounts (yellow). The reversal points at the cue were estimated using the linear regression depicted in (A)*. ***(C)*** *Decoded joint density of reward over magnitude and time, using the population responses at the variable and certain cues, and the single neuron estimated tuning functions for reward magnitude and time*.

**Figure 5:**
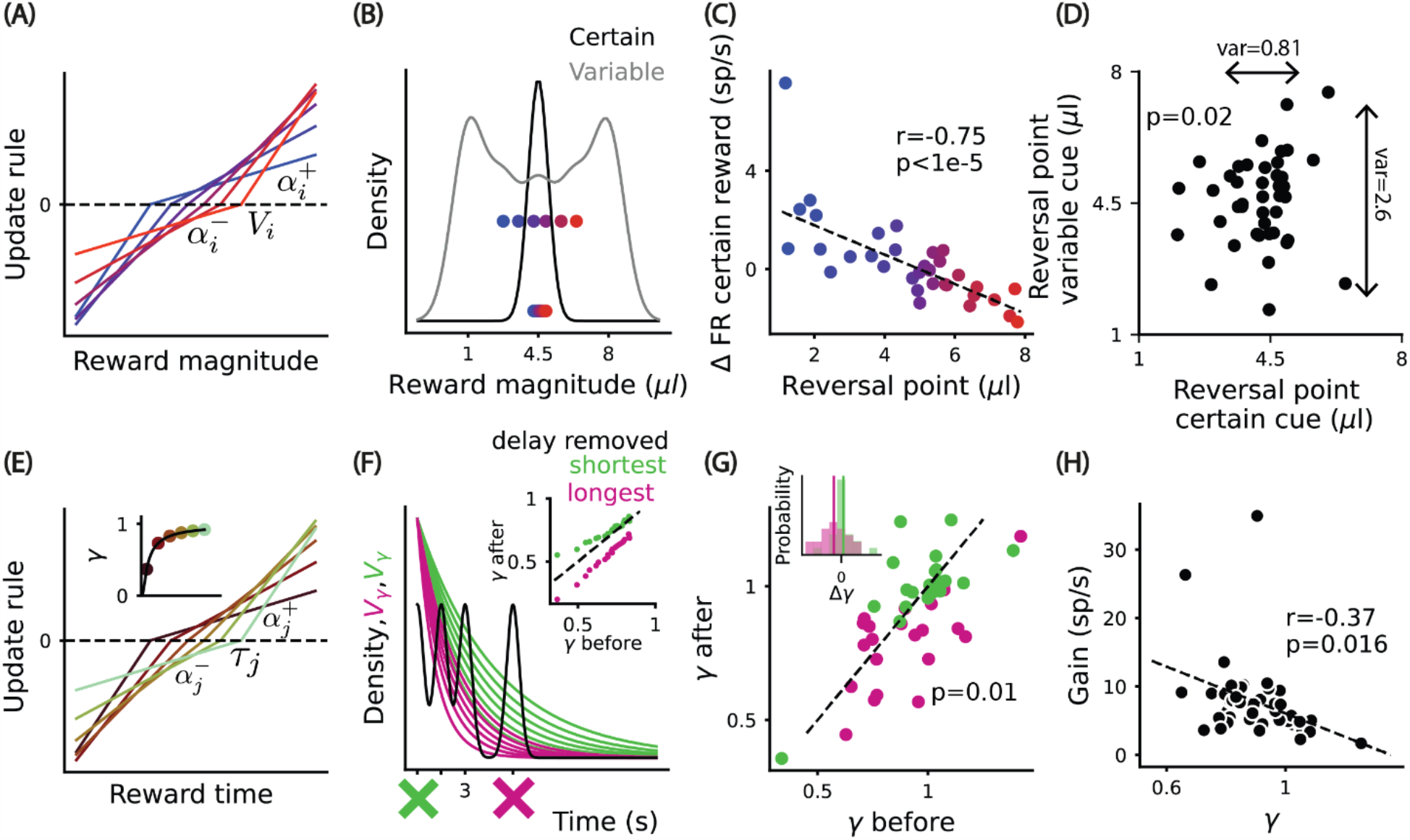
Value and temporal sensitivity adapt to changes in reward statistics, in accordance with principles of efficient coding. ***(A)*** *The distributional value code predicts that units have different asymmetries for positive* 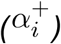 *and negative* 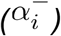 *RPEs, generating a set of values V*_*i*_, *color code: reversal point*. ***(B)*** *The distributional value code predicts that variability in reversal points for the cue predicting a bimodal distribution is greater than for the cue that predicts a certain reward amount*. ***(C)*** *DAN responses to the certain reward amount delivered at a 3s delay, as a function of the reversal point estimated using the responses for the variable reward amounts delivered at a delay of 3s. P*earson correlation coefficient r=-0.75, two-tailed test p-value<1e-5, 95% CI=(-0.*87,-0*.*54)*. ***(D)*** *Reversal points estimated at the cue predicting variable and certain reward magnitudes at the same delay. The reversals were estimated using the firing rate at the cue corrected for the diversity in temporal discounting and the linear regression depicted in Figure 4A. The variance is depicted alongside double-ended arrows. Bootstrapping 10000 times to test if variance of reversal points for variable cue is greater than for certain cue: p-value=0*.*02, mean difference in the variances=1*.*71, 95% CI=(0*.*15,6*.*9)*. ***(E)*** *Considering asymmetric weights for under and over-estimation of reward times generates a diversity of time constants that are mapped to temporal discount factors using the function depicted in the inset, color code: temporal discount factor*. ***(F)*** *Predicted adaptation in temporally discounted values when the reward time distribution is manipulated, by removing the shortest (green curves) or longest delay (magenta curves). The black curve depicts the smoothed distribution of reward times in the first phase of each experimental session. The inset depicts predicted adaptation in temporal discount factors*. ***(G)*** *Experimentally observed adaptation in temporal discount factor estimated from the recorded DANs. Responses to the same delays were used to estimate the temporal discounts before and after the manipulation. The inset depicts the histogram of the update in temporal discounts for the two different manipulations. The vertical lines depict the mean update. Bootstrapping 10,000 times testing if update in temporal discount when removing the shortest or longest delay is different: p-value=0*.*01, mean absolute difference=0*.*13, 95% CI=(0*.*008, 0*.*27). Bootstrapping 10,000 times testing if update when taking longest delay is negative:* p=0.04, 95% CI=(-0.34,-0.0070). *Bootstrapping 10,000 times testing if update when taking the shortest delay is positive:* p-value=0.3, 95% CI=(-0.07,0.22). ***(H)*** *Single neuron gains as a function of temporal discount factors, the dashed line represents the fitted linear regression*. Pearson correlation coefficient r=-0.37, two-tailed test p-value=0.02.

A key prediction of our distributional code for reward timing is that the probability distribution over future reward time can be decoded from the population responses to a CS (Figure 1C). We focus on the population responses at the CSs that predict a certain reward at fixed delays and assume the system has knowledge of the temporal discount rate of each neuron. Since the population response at the CS reveal the encoded values, assumed to reflect the sum of temporally discounted rewards over time, determining the distribution over future reward time presents as a linear regression problem (Figure 3D top). The independent variable is a matrix of temporal discounts to the power of the discretized time, the dependent variable is the mean responses at the CS and the regression coefficients are the probabilities of rewards over time (Figure 3D top). The resultant densities capture the differences in the timing of rewards for the four CSs (Figure 3D bottom).

We next asked if the decoded estimates correspond with animals’ temporal expectations, by comparing trial-by-trial variability in anticipatory licking behavior with the future reward time predicted by the population of DANs. We observed that in trials wherein animals commenced licking earlier or later, the decoded distribution over future rewards exhibited a qualitatively similar shift in time (Figure 3E). These data suggest that estimates of future reward times decoded from a dopamine neuron population reflected temporal expectations that animals used to guide behavior.

### Dopamine neuron cue responses reflect distributional value information encoded in responses to reward

Next, we sought to combine the distributional time code with the previously proposed distributional code in amount^7^, focusing first on the CS predicting a variable reward amount. The distributional RL theory in amount predicts that neurons with asymmetric linear functions for positive and negative RPEs possess reversal points (the reward amount for each neuron that produces zero net change in activity) that correspond to the expectiles of the probability distribution of rewards^7^. Previous work linking distributional codes for reward to the activity of midbrain dopamine neurons has largely focused on reward responses, and not the responses to cues that predict rewards. However, in principle, distributional information should be propagated backward in time from reward and thus decodable from the responses to reward predictive cues (Figure 1D). One reason this may not have been observed in previously reported data is that the diversity in temporal discounting across neurons that we describe in the previous section can occlude distributional reward magnitude information (Figure 1A,D). Because we measured diversity in temporal discounting across neurons, we were able to correct for it. Indeed, we found that correcting for the diversity in temporal discounts and gains revealed residual CS responses that are significantly correlated with the reversal points estimated at the time of rewards (Figure 4A), indicating stable but mixed selectivity for reward delay and magnitude in single neurons. We then computed probability distributions over reward magnitude from dopamine responses at both the time of CS and the time of reward delivery. The distributions from reward and cue responses were nearly identical (Figure 4B), indicating that the consequences of systematic variability in sensitivity to positive and negative of errors at reward delivery, previously identified as evidence that the dopaminergic system implements a distributional code for value, are transmitted to the cue response. This establishes the presence, at the time of cue presentation, of information required for the system to foresee impending variability in the magnitude of rewards ahead of time, when it might be used to guide future behavior.

We then tested if the joint distribution over future reward amounts and times can be decoded, a key prediction of the distributional TMRL code, from the responses of the DAN population at the CS. Conditioned on the CS, the joint probability distribution of reward on each trial can be factorized as the product of the marginal distributions of reward over time, giving a 2D map of future reward magnitude over time (Figure 4C) that closely matches the true distributions. Thus, a multi-dimensional probabilistic map of future rewards may be estimated from just 450ms of dopaminergic neural activity at the onset of an episode.

### Value and temporal sensitivity efficiently adapt to environment statistics

We have identified diverse temporal discounting and sensitivity to value in midbrain DANs that may be used to estimate the joint probability of future reward time and magnitude. However, does parameter diversity across neurons that enables such a code reflect variability that we as experimenters exploit? Or are the parameters collectively regulated to maximize information about rewards along the hypothesized target dimensions? Evidence of such regulation would not only indicate efficient representational codes for reward within the dopamine system, but by extension, also provide strong evidence that the diversity in parameter tuning is present for the purpose of representing distributional reward information.

We used an efficient population coding framework^22–24^ to derive temporal discount functions that optimally represent the reward times *t*_*r*_ in the environment. Inspired by previous work, we propose that this distribution is efficiently encoded in an expectile code^7^. In particular, we sought to maximize the *mutual information* between the true expectile reward times predicted at the cue *s*,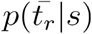, and those encoded by the dopamine neural population 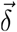, constrained by the number of neurons *N* and by the population expected firing rate *R*. We assume the cue response of each dopamine neuron decays exponentially as a function of reward delay with a time scale *τ* and a gain parameter *a*. Parameterising the population with the density of tuning curves *d* and a gain *g*, the solution is that neuron’s timescales *τ* should distribute according to the probability distribution of the current environment expectile reward times 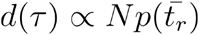 (see Methods and Ext. Data Fig. 10). Intuitively, if rewards at short delays occur infrequently in a given environment, the system should not waste coding capacity to encode expected future rewards at short timescales, and vice versa if the environment only rarely emits reliable rewards at long delays. Furthermore, for each neuron the gain should be inversely proportional to the probability that a randomly chosen reward time will be smaller than it’s time scale *P* (*τ*), *g*(*t*_*r*_) = *R*/*NP (τ*)), (see Methods). For low reward times, the entire population is active, incurring a large metabolic cost for encoding these values. Intuitively, this metabolic penalty can be reduced by lowering the gains tuned to late reward time scales, while maintaining optimized coding.

In order to optimize this efficient population code online, we generalize the distributional learning rules (Figure 5A^6^) to the time domain, considering multiple channels with different relative scaling for over and underestimation of reward times, that generate a diversity of learnt reward time scales (Figure 5E),

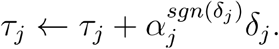

Importantly, these parameters converge to the efficient code that optimally adapts to the statistics of expected reward times in the environment (see Methods).

A critical but untested prediction of the distributional code for value is that the value each neuron expects (as defined by its reversal point) should adapt to changes in the probability distribution of reward magnitudes (Figure 5A,B). In addition, the ordering of reversal points in the population should be preserved for different probability distributions. We found that the reversal point ordering is preserved across the two CSs (Figure 5C) and that the variance of the reversal points for the variable CS is significantly greater than for the certain CS (Figure 5D).

The mapping from time scales to temporal discount factors is an exponential mapping that takes into account the fact that steep temporal discounting (ie. small temporal discount factors) gives rise to very different learned value estimates for rewards that will occur at distinct short delays, but discriminates poorly between rewards occuring at later delays. Conversely, shallow temporal discounting (ie. large temporal discount factors) discriminate between rewards occurring far apart in time, and can thus support encoding of rewards at long time delays. Therefore, the discount factor is a monotonically increasing, exponentially decelerating function of the time scales at which rewards are observed 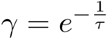 (Figure 5E inset).

To test if the dopamine code adapts to the temporal statistics of reward, at the end of each session we modified the temporal reward distributions by removing the CS corresponding to either the shortest or longest reward delay. As predicted by the theory (Figure 5F), when removing the longest delay, DANs adapted to improve encoding accuracy on short reward times by decreasing their discount factor (Figure 5G). While adaptation when removing the shortest delay was not statistically significant in this dataset (Figure 5G), the magnitude of the theoretically predicted adaptation is smaller than that predicted when removing the longest delay, and the data exhibited a trend in the correct direction, and thus a larger data set may reveal a shift in this condition as well. Alternatively, asymmetry in neural adaptation may be due to a rational bias in the neural code towards ensuring that predictions for the very near future are accurately encoded regardless of the temporal reward distribution in the environment, given that passage through intervening delays en route to later rewards is unavoidable. We also tested whether discounting by DANs was updated more continuously by comparing the trial-to-trial adaptation in temporal discount when the rate of reward occurrence is relatively high or low. Indeed, the temporal discounts for low rates were larger than for high rates (Ext. Data Fig. 9). Regarding the gain, we indeed observe as predicted that individual neuron gains are negatively correlated to temporal discount factors (Figure 5H).

The lawful, dynamic regulation of temporal discounting in individual DANs that we describe here indicates that principles of efficient coding likely apply to how the brain regulates the time constants over which rewards are predicted. However such lawful adaptation also strengthens our confidence that the dimension over which we as experimenters are decoding expected rewards - time - is indeed an encoding target for the system.

### How might information about future reward distribution be used to guide behavior?

The TMRL algorithm is relatively simple and “model-free”, meaning that the algorithm itself does not have access to knowledge of how the world transitions between states when learning to assign value to them. However, we reasoned that the very nature of the information it learns can naturally allow for complex computations such as planning. To examine the potential behavioral benefits of the joint distributional reward representation produced by TMRL in relation to pre-existing algorithms, we performed a series of simulations in which model agents forage for rewards (Figure 6). For comparison, we model agents making use of either TMRL, standard TDRL, or the successor representation (SR). Similar to TMRL and TDRL, SR is a predictive model learned using a temporal-difference algorithm. However, SR instead caches expectations of future state visits, which can facilitate heuristic planning strategies^25^.

**Figure 6:**
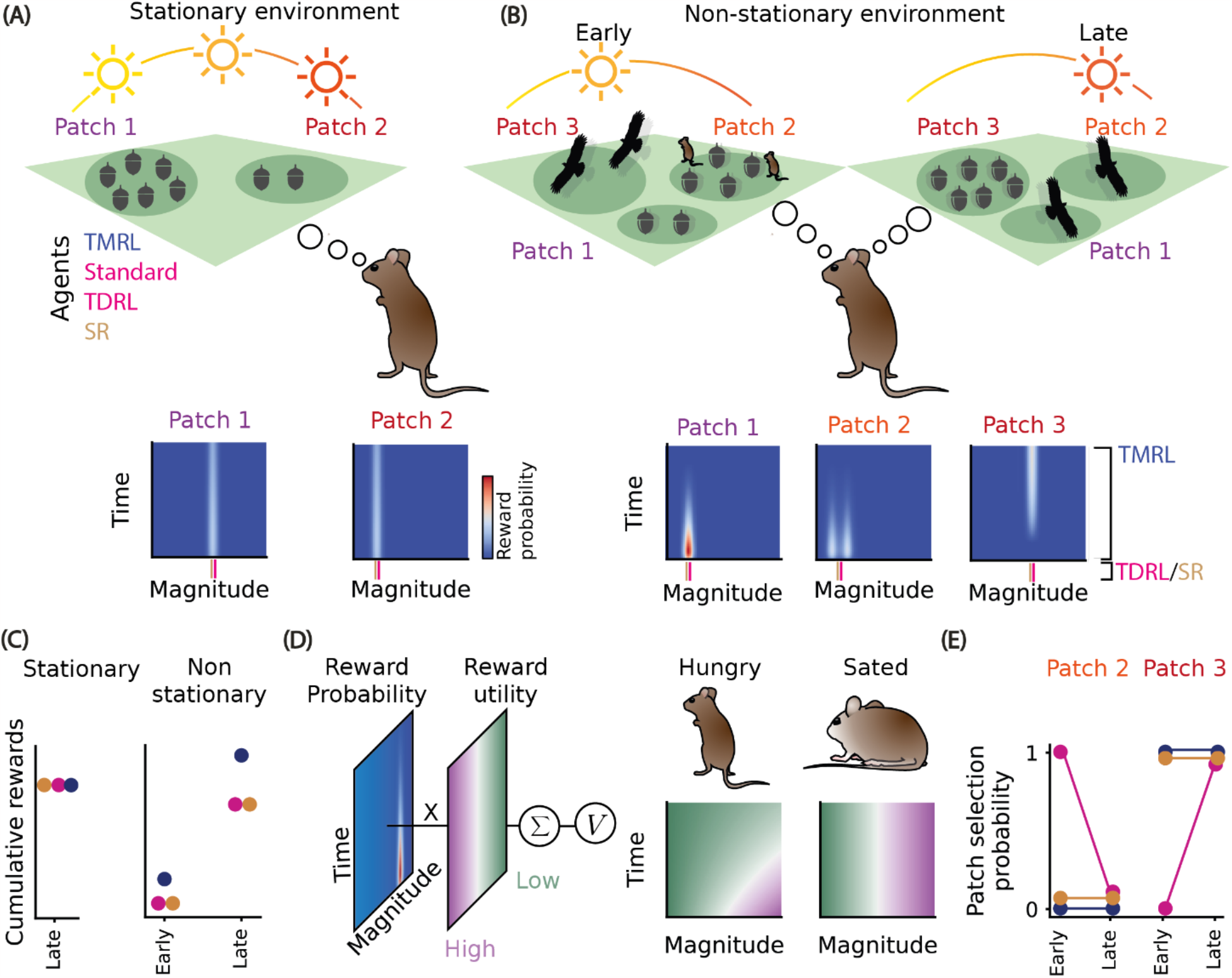
A distributional code allows for planning and flexible adaptation to temporal dynamics and preferences of reward using a model-free RL algorithm. ***(A)*** *A foraging mouse must decide which patch to choose to maximize cumulative collected rewards in a stationary environment. Axes indicate the learned joint probability distribution of reward time and magnitude associated with each patch. Rewards available from the two patches are stable during the day, with patch two providing a larger quantity of acorns than patch one*. ***(B)*** *In the non-stationary environment patch one provides fewer acorns early in the day. Patch two provides a variable (represented as competitors) number of acorns with the same mean as patch one early in the day. Patch three provides more acorns than either patch one or patch two, but later in the day. The reward variability in magnitude can arise due to the presence of competitors (represented as other mice). On the other hand, temporal dynamics can arise from environmental factors such as the foraging strategies of predators (represented as eagles, which preclude acorn availability). The standard TDRL and SR only keep track of a scalar value V (represented as vertical lines below the axes), whereas the TMRL keeps track of the future probability over reward time and magnitude*. ***(C) Left:*** *Cumulative rewards obtained when simulating the behavior of the three different algorithms in the stationary environment*. ***Right:*** *Cumulative rewards obtained when simulating behavior of the three algorithms in the non-stationary environment when fewer (early in learning) or greater (late in learning) numbers of time steps have elapsed*. ***(D)*** *In the agent making use of TMRL, the probability distribution over future reward time and magnitude is additionally weighted by a utility function to obtain an estimate that depends on internal state (see Methods for a detailed description). The utility function when the mouse is hungry (center) or sated (right)*. ***(E)*** *Probability of choosing patch two (left) and patch three (right) when the mouse becomes sated early or late in learning, color coded for the three different algorithms*.

Interestingly, the basic mechanism underlying the SR - TD learning of expected future state occupancy - has recently been extended to include a set of SRs that vary in their temporal horizon, a conceptually similar innovation to that reflected in TMRL, except that it focuses on creating multi-scale temporal predictions of future states instead of future rewards^26^.

Our simulations took the following form. In a simulated stationary environment where a mouse can choose between patches that deliver different but stable amounts of reward (eg. acorns), the TDRL, SR and TMRL agents perform similarly (Figure 6A,C). However, in a dynamic environment where different patches have different amounts of available reward at different periods of time during the day, mice can collect significantly more reward when given access to knowledge of the timing of reward availability (in addition to the average amount of reward each patch contains). In naturalistic settings, dynamics of effective reward availability may arise due to resource production itself, foraging strategies of competitors, or dynamic predation^27^. By forward planning over the map generated by the distributional TMRL algorithm, we find that mice can adopt a strategy where they harvest from a lower value, but more immediate, reward patch (patch one and two) before moving on to a higher value, but delayed, patch (patch three) that would be persistently selected by a single temporal discount factor model that only learns about the mean reward of each patch (Figure 6B,C). The TMRL also outperforms the SR agent, since the standard SR algorithm also estimates a scalar value. These simulations provide a simple demonstration of the potential benefit of using information about the distribution of rewards in time to guide action selection over existing models that do not possess such information, specifically in the context of an environment with temporally delimited reward availability.

The previous example focused on a case where the external world possesses reward dynamics, but biological agents are also subject to internal state dynamics that incentivise using knowledge about reward distributions. For example, in response to changes in wealth or physiological need state, humans and other animals shift their preferences with respect to the delay and variability in the amount of reward, which can be expressed as a utility function^28–30^,^31,32^. The distributional TMRL code generates a representation that may be used to flexibly adapt policies to these preferences, as has been shown previously for model based implementations^33^. For example, if the mouse is hungry and urgently needs a significant amount of reward, they might reweight different regions of the TMRL reward map, akin to creating a dynamic utility function that heavily weights large, imminent rewards (Figure 6D). This might lead the mouse to employ a policy where they select patch two, predicted to grant the largest reward at a short delay. When sated, the mouse may want to devise a different utility function that does not vary as a function of predicted reward delay, initially selecting patch three, which gives the largest reward at the longest delay (Figure 6D). The distributional TMRL algorithm and the SR allow for a faster adaptation of policies from the hungry to the sated state than the TDRL algorithm, which needs to learn through experience to adapt the policy (Figure 6E), and thus may present interesting potential computational mechanisms for how internal states modulate decision-making.

While not intended as exhaustive, the two scenarios we simulate here in a foraging environment - temporally delimited and variable magnitude rewards, and internal state dependent modulation of utility - are chosen to illustrate the broad potential importance of the information provided by distributional TMRL for animal behavior. They reflect a small, but illustrative, sample of the rich opportunities for future work to examine how the reward representations generated by a multidimensional distributional algorithm like TMRL can benefit adaptive behavior across a range of scenarios.

## Discussion

A fundamental facet of intelligence is the ability to use past experience to predict the future^34,35^. Predictions may incorporate detailed information about how the environment will develop depending on a course of action, constituting models of the world that can be operated on flexibly but laboriously. Alternatively, predictions may take the form of efficient, compressed representations that discard detailed features of environmental structure and are specific to particular, behaviorally relevant events, such as rewards. Within RL, model-based learning algorithms that involve the former, detailed predictions enable more flexible behavior at the cost of computational complexity, while model-free algorithms that target the latter, simpler predictions allow for efficiency at the expense of flexibility. The field of RL provides a growing set of tools for learning simpler and more complex predictions alike in service of adaptive behavior, and there is ample evidence that the brain employs strategies resembling both algorithmic classes depending on, for example, whether behavior is under more explicit, goal-directed or automatic, habitual control^33,36^.

Though often described in categorical terms, recent work has highlighted how predictive representations of intermediate complexity can enable more flexible behavior that tends to characterize model-based algorithms, while using computationally efficient model-free learning algorithms^25,37^. Here we present distributional TMRL, a theoretical extension of TD learning that learns an efficient, multi-dimensional, probabilistic map of future reward value over time. This probabilistic map constitutes a kind of ‘model’ of the world, and yet, the mechanics by which it is learned are less complex than those used to learn the full structure of possible state transitions in an environment. Furthermore, we show how distributional TMRL learns representations that may enable better, ie. more rewarding, policies for behavioral control when in an environment where exploiting knowledge about the time course of available rewards confers a benefit, or when changes in internal state might favor dynamic attitudes toward risk. Importantly, we present evidence that the brain may make use of TMRL-like computations. Midbrain DANs, a core component of neural systems involved in learning policies for behavioral control, displayed signatures of a multidimensional distributional code over reward time and value; furthermore, animals appear to use the reward timing information reflected in DAN population activity to guide temporal control of behavior. Lastly, we show that the distributional TMRL code adapts to changes in reward statistics in accordance with information theoretic principles of efficient coding. This provides not only strong evidence that the joint distribution of reward over time and value that we as experimenters decode from neural activity is indeed a functional target for encoding within the system, but potentially opens a fresh perspective on how to interpret individual variability in temporal discounting at a behavioral level. Steep temporal discounting that heavily favors immediate over delayed rewards, leading to apparently maladaptive, impulsive behavior, can be reframed as the optimal solution to a volatile environment. Conversely, shallow discounting, leading to consideration of delayed rewards, is ideal for stable environments that are predictable over long time scales. Thus, changes in environmental volatility may be compensated by adaptive changes in discounting, complementing evidence for volatility-induced changes in learning rates^38–40^. This suggests that therapeutic attempts to improve impulse control might fruitfully target the environment, by intervening to lengthen the timescale over which predictions are valid, or reorient individuals to the longer timescale structure of rewards that may already exist within an environment.

More than two decades ago, midbrain DANs were first hypothesized to emit a TD RPE to teach recipient circuits accurate reward expectations^1,5^. This proposal drove the development of a new field of research into whether the brain may be using algorithms like those contained within computational RL models to learn programs for behavioral control, and if so, what form they take. That a compact algorithm like TD learning can estimate an overarching objective around which much of learned behavior can be shaped is not only practically useful in engineering settings, but explains a wide range of neural and behavioral data. However, in its simplest form, TD learning does not capture behaviorally relevant dimensions that characterize rewards because it compresses information regarding reward time, magnitude, and quality into one scalar value that represents, in a common currency, the expected average sum of temporally discounted future rewards. Sensory characteristics, and unambiguous knowledge about magnitude and timing of rewards is lost. Furthermore, experimental data has shown that DANs can exhibit sensitivity to the sensory characteristics of rewards ^41^, actions or features of the environment^42^, and appear to access internal models regarding environmental structure^43^. It is thus becoming ever more clear that the basic model of DANs emitting a unitary, model free TD RPE requires refinement, at the very least, and some have argued for its replacement altogether^44,45^.

However, the space of possible TD learning algorithms is large and continually growing. Assumptions about the way the state of the environment is encoded, the space of possible actions in TD methods for control, and parameter determination or regulation are just a few of many ways in which TD methods can differ ^46^. Given the parallel and modular nature of brain circuits, including those embedding DANs, one class of TDRL extensions that would seem particularly attractive for deriving hypotheses about neural systems posits parallel learning of multiple reward expectations that systematically differ, either quantitatively or qualitatively. For example, multi-system models, where separate model-free and model-based RL mechanisms both contribute to behavioral control have been mapped to distinct, parallel circuitry in the basal ganglia ^47,48^, the main target of dopaminergic innervation, and multi-agent and mixture of experts models have been used to explain recent data regarding the functional role of neurons in different regions of the striatum, a major target of dopamine, and spatial heterogeneity in striatal dopamine signaling itself, respectively^49,50^. In addition, recently described heterogeneity in DAN encoding of task variables has been hypothesized to reflect a vector, as opposed to a scalar, prediction error, still within a TD framework ^51^, and signatures of multiple separate, qualitatively distinct predictions have been observed in DANs^52^. Here we provide direct evidence that diverse sensitivity to a fundamental dimension along which rewards are distributed, time, can explain another source of variability in response properties across DANs, without abandoning a computational framework for understanding dopaminergic function for which the accumulated experimental support is large.

Temporal information is critical for learning systems. Reliable temporal structure in the world forms the basis for identifying predictive relationships that drive associative learning and contribute to causal inference. To extract such structure, the brain must somehow register when certain events occur in relation to each other, creating something akin to a temporal map^53^. We describe the existence of just such a temporal map, specific to reward, that is reflected in a core component of reward circuitry long known to be critical for behavioral control: midbrain DANs. It will be important to determine in future work whether and in what manner this information is distributed within other brain circuits, and how this information is brought to bear on the problems faced by organisms seeking to thrive within complex and dynamic natural environments.

## Acknowledgements

We thank H. Schuett, F. Rodrigues, T. Duarte and C. Haimerl for comments on versions of the manuscript and the entire Paton laboratory, past and present, for feedback during the course of this project. The work was funded by an HHMI International Research Scholar Award to J.J.P. (55008745), a European Research Council Consolidator grant (DYCOCIRC - REP-772339-1) to J.J.P., a Bial bursary for scientific research to J.J.P. (193/2016), internal support from the Champalimaud Foundation, and PhD fellowships from FCT to M.S. (PD/BD/141552/2018) and B.F.C. (PD/BD/105945/2014). The funders had no role in study design, data collection and analysis, decision to publish or preparation of the manuscript.

## Author contributions

M.S. developed the theory, together with J.J.P and D.M and with input from K.L. Experiments were designed by J.J.P, M.S. and B.F.C. The experimental apparatus was constructed by. B.F.C., P.B. and M.S. All behavioral and electrophysiological experiments that provided data for the study were performed by P.B., and B.F.C. performed pilot experiments to establish protocols. P.B. and M.S. analysed histological data. M.S. analysed the neural and behavioral data, with input from B.F.C., K.L., and D.M. M.S. performed the foraging simulations with input from D.M and K.L. J.J.P. and M.S. wrote the paper, and D.M., K.L., and B.F.C. edited the paper. J.J.P. supervised all aspects of the project.

## Data Availability

All data generated or analyzed during this study (and its supplementary information files) will be made available upon publication.

## Code Availability

The analysis code and code used to generate the theoretical predictions and simulations will be published in an online repository.

## Methods

### 1 Mice

Young adult (2-7 months old at time of experiments), male DAT-Cre mice expressing Channelrhodopsin-2 (ChR2) in midbrain dopamine cells were used in this study. For this, Ai32(RCL–ChR2(H134R)/EYFP) mice (IMSR_*JAX*_ : 012569) were crossed with DAT-IRES-Cre mice (IMSR_*JAX*_ : 006660). Mice were group-housed (up to 3 mice per cage) until the first of two craniotomies were performed, after which they were single-housed. A temperature (21°C) and humidity-controlled (50%) housing room was maintained with a 12-hour light/dark cycle. Mice were maintained on PicoLab Rodent Diet 20 (5053), and under water deprivation for all behavioural experiments (*>* 85% body weight from baseline *ad* libitum period before deprivation). All experiments and procedures followed guidelines set and approved by the relevant national and international authorities (Champalimaud Foundation Animal Welfare Body (protocol number: 2017/013), Portuguese Veterinary General Board (Direcção-Geral de Veterinária, project approval 0421/000/000/2018) and European Union Directive 2010/63/EEC).

### 2. Surgical Procedures

Each mouse received three surgeries, first one for headpost implantation and then two unilateral craniotomies (performed 1 week apart) above the targeted regions (VTA/SNc). All surgical procedures were carried out under anesthesia with isoflurane (3% for induction; 1–2% for surgery at 0.8 l min ^−1^). Mice were then fixed in a stereotaxic frame and their eyes were protected with a small amount of ophthalmic ointment.

The headpost implantation surgery was performed following the procedure outlined in^61^. Briefly, the hair was shaved down to skin which was then disinfected and incised. After exposing the skull, the soft tissue and periosteum were carefully removed and cleared. Then bregma and lambda were identified, the skull was leveled, and the locations for the craniotomies were marked. Subsequently, the skull surface was prepared for headpost implantation following the procedure suggested in the protocol. The headpost was then placed in a desired location (anterior to bregma and slightly above the skull surface) and cemented in place with dental adhesive (C&B Metabond, Parkell). Following the surgery, mice received carprofen subcutaneously (s.c.) for pain management. Mice were then placed back in their original cages and monitored for 7 days post-surgery to ensure well-being and a full recovery.

Prior to any electrophysical recordings (24-48 hours), a small craniotomy (1.5mm) was performed and sealed with a removable silicone sealant (Kwik-Sil, World Precision Instrument). Two separate craniotomies, one in the left and the other in the right parietal bones, were performed (coordinates: AP: -3.0 mm; ML: ±0.6 mm from the Bregma). Prior to surgery, all mice received carprofen (s.c.), enrofloxacin (s.c.), and dexamethasone (i.m.).

### 3 Behaviour & Training

#### 3.1 Behavioural apparatus

The behavioral setup consisted of an infrared light source, an IR camera, a head-fixation system, a water delivery tube, a custom-built olfactometer^62^ with an odor delivery tube, and an air ventilation system that was running during the whole experiment to prevent odor accumulation. The task logic was implemented in a real-time operating system using an Arduino microcontroller (Arduino Mega 2560, Arduino). The behavior of the animal was also monitored via an IR camera (FL3-U3-13S2, FLIR). The videos were acquired with Bonsai^63^ at 120fps (640 x 480 pixels) for online licking detection and further offline processing. Briefly, a small area of the image was selected, on each session. For each frame, the sum of all pixels’ luminance was computed. The resulting trace was then manually thresholded and lick events were detected as frames with average luminance above the background.

The odor cues were delivered through the tube of a custom-built olfactometer placed approximately 1cm from the mouse snout, and the odor delivery was controlled via two-way micro-solenoid valves (model LHDA1233115H, Lee Company, CT, USA). Similarly, calibrated water reward was delivered through a lick spout using a two-way micro-solenoid valve.

#### 3.2 Odor stimuli

During each trial an odor cue was presented for 1 second approximately 1 cm from the snout. Odors were delivered via a computer-controlled olfactometer with a 1,000 ml/minute constant flow. Each odor was dissolved in mineral oil at 1:10 dilution and 15*μ*l of diluted odor solution was applied to the syringe filter (2.7*μ* m pore, 13mm; Whatman, 6823-1327). Odors were: Cuminaldehyde, (S)-(+)-2-Octanol, (R)-(-)-Carvone, Pentyl acetate and Hexanoic acid.

#### 3.3 Behavioural task

Mice were water restricted 7 days after head-posting and habituated to head-restrainment for 2-3 days (10-30 min sessions). Within these sessions, mice were allowed to voluntarily lick the spout for a water reward. Following the habituation sessions, mice were trained in an odor-cued classical conditioning task, where an odor cue predicts reward with distinct delay and/or magnitude (*i*.*e*. volume of water).

The task included five trial types, randomly intermixed. Trial types 1-4 began with a 1-s odor delivery, followed by a delay of 0, 1.5, 3, 6 seconds, respectively, and a fixed water reward amount of 4.5 microliters. For one of the animals, the delay of 0s wasn’t included in the task. Trial type 5 began with a 1s odor delivery, followed by a 3 seconds delay and a reward sampled from a probability distribution with five possible outcomes: 1, 2.75, 4.5, 6.25, 8 microliters with respective probabilities: 0.25, 0.167, 0.167, 0.167, 0.25, such that the mean of this distribution was 4.5 microliters and thus matching the average reward amount delivery across all trial-types. The odor identity associated with each trial type was shuffled for different individual mice. Importantly, at the end of the session, when 200 trials had passed, there was a context switch and either the cue predicting the shortest or the longest delay was removed. Trial duration was drawn from an exponential distribution (minimum 11s, mean 4.5s, truncated at 21s), resulting in an approximately flat hazard function and an approximately constant reward rate throughout the session.

### 4 Electrophysiology

#### 4.1 Acute recordings

All electrophysiological experiments were conducted while mice were head-restrained. Recordings were performed for up to 6 days following a 24-48 hours recovery period from the craniotomy surgery. Between recording sessions, the craniotomies were covered in the same manner as described above. A two-shank 64-channel silicon probe (ASSY 77-H6, Cambridge NeuroTech) with a tapered optical fibre (Lambda-B fibre 100-μm core NA=0.48, Cambridge NeuroTech) glued to the back, was lowered into the ventral tegmental area (VTA) and substancia nigra pars compacta (SNc). Probe tracks were reconstructed from three different sessions by dipping the probes in DiD, DiO and DiI solutions before insertion. Electrophysiology and laser/LED modulation data were digitized at 30kHz with the Open Ephys acquisition board^64^ and recorded with Bonsai^63^.

### 4.2 Spike sorting and data processing

In order to remove light artifacts, independent component analysis (ICA) was performed^65^. In particular, we used chuncks of recorded signals from the beginning and end of each session where pulses of light were given, and used fastICA^66^ to obtain the independent components and the mixing matrix. Light artifacts were present simultaneously in the majority of the channels, hence we considered as artifacts the two components with highest entropy of the mixing matrix weights, and after visual inspection, removed these components and reconstructed the signals. Data were sorted offline with Kilosort 2.5 spike sorting software (https://github.com/MouseLand/Kilosort), and manually curated using Phy (https://github.com/cortex-lab/phy).

### 4.3 Light identification of dopamine neurons

The light evoked photoactivation of midbrain dopaminergic neurons with ChR2 was used to identify neurons as dopaminergic. The protocol consisted of trains of 10 blue (473 nm) light pulses, each 10 ms long, at 1, 5 and 20 Hz (5s inter-train interval) followed by 3 consecutive long pulses, lasting for 1s each. Laser power at the tip of the fiber was on average 20 mW (measured at the tip of the fiber, before each experiment). Optogenetic stimulation was delivered twice at the beginning and twice at the end of the recording session. Units were considered photo-identified using an intersection of different criteria: SALT test^67^ p-value *<*0.001, paired t-test comparing baseline and post-laser onset (1–10ms) firing rate yielding a p-value*<*0.01, correlation coefficient between laser-triggered waveform, non-evoked waveform *>*0.9 and probability of eliciting ≥1 spikes within 1-10ms of each pulse *>*0.1. To be included in the data set, a neuron had to be well-isolated (inter-spike-interval (ISI) violation *<*0.04^68^).

### 5 Immunohistochemistry and microscopy

Histology was performed to verify placement of the recording electrodes and expression patterns of transgenes. The mice received an overdose of pentobarbital (Eutasil, 100 mgkg^−1^ intraperitoneally) and, once deeply anesthetized, were perfused transcardially with 4% paraformaldehyde. The brains were removed from the skull, stored for 24h in 4% paraformaldehyde and then kept in PBS until sectioning. Coronal brain slices were obtained using vibratome sectioning (80 *μ*m), and immunostained with antibodies against GFP: rabbit anti-GFP (A-6455, 1:1,000) and goat antirabbit AF488 (Invitrogen, A-11008, 1:1,000). The sections were incubated in DAPI and mounted In Mowiol. Imaging was carried out using a slide scanner (Axio Scan Z1, Zeiss). For electrophysiological recordings, shank placement was confirmed using DiD, DiO and DiI cell-labeling solutions (V22889, Thermo Fisher Scientific).

### 6 Probe trajectories & location of the recording sites

Probe trajectories were reconstructed from histology data by alligning the histological slices with the Allen common coordinate framework (CCF) atlas and manually tracing the dye track using code that is publicly available at https://github.com/petersaj/AP_histology.

CCF coordinates were transformed into stereotaxic coordinates using the method described in https://community.brain-map.org/t/how-to-transform-ccf-x-y-z-coordinates-into-stereotactic-coordinates/1858. The AP, ML and DV coordinates were estimated for the probe’s tip (visual guess) and 400 points that lie within 400 μm (step k = 1 μm) from the tip. In total, 54 cells (18 photo-identified and 36 putative) were recorded from the sessions with the dyed probe. The coordinates were estimated for the channels these cells were recorded from with the depth of each cell taken from the Kilosort results.

### 7 Distributional code for reward time model

We describe our model in the context of a class of Markov decision processes (MDPs) which use a deterministic chain of states to model the temporal evolution of a reward function across time. This is known as the complete serial compound representation^69^. Such MDPs (𝒯, 𝒫, *r*) are composed of a set of time states 𝒯, a reward random variable *r* : 𝒯 → ℝ indexed by time states and a deterministic transition function between time states 𝒫 (*t* + 1|*t*)= 1. The value at time step *t* is the expected sum of discounted future rewards over a time range from 0 to *T*,

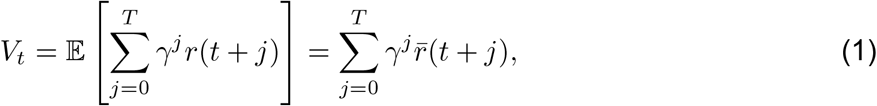

where 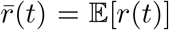 and *γ* ∈ [0, 1] is the temporal discount factor that determines how much the utility of delayed rewards is decreased relative to immediate reward.

#### 7.1 Efficient population coding and the expectiles of reward times

In our experiment, reward time and amount is manipulated across conditions and thus our dopamine population model describes adaptive neural responses to both these features of the environment reward function. Initially, we describe a dopamine neuron’s tuning function with respect to reward time *t*_*r*_. It is proposed that, across the dopamine neuron population, reward is heterogeneously discounted over time, and reflect this assumption in exponentially decaying neural tuning function parametrized by reward time scale *τ*, with a gain parameter *a*

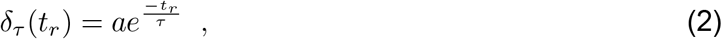

where *τ* is the time at which the neural response is reduced to 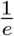 times its initial value. Considering this parametrization, the corresponding temporal discount factors (see Eqn. 1) arise naturally as a function of the reward time scales, specifically 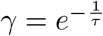. We consider a mapping *ι* : 𝒯 → *T* from the set of neuron’s reward time constants 𝒯, to the space of reward times *T* .

In order to predict optimal variability across the reward time scales estimated from dopamine neuron responses, we develop an efficient population coding model^22^ for the distribution *p*(*t*_*r*_|*s*) := *p*(*t* = *t*_*r*_|*r* ≠ 0, *s*) of reward times *t*_*r*_, where *s* is the stimulus presentation at time 0. That is, the distribution of possible future reward times *t*_*r*_ after the stimulus has been observed. We propose that this distribution is efficiently encoded in an expectile code, as opposed to the quantile code. Indeed, previous experimental work has shown that the tuning functions of dopamine neurons of reward magnitudes are more consistent with bi-linear functions, that would be predicted for an expectile code, then with heaviside functions, that would be predicted for a quantile code^7^. Thus, for consistency with these previous results, we use an expectile code model based on expectations of reward times. Theoretically, we accomplish this by optimizing our dopamine population model to efficiently encode the distribution of expectations over reward times. In contrast, the quantile code may be computed in our formalism by efficiently encoding the distribution over reward times^70^. Importantly, though our theoretical formalism is general and integrative over both expectile and quantile codes, our use of the expectile code in particular does not qualitatively change our model predictions.

The expectiles of a distribution generalize the mean statistic analogously to how quantiles generalise the median^70^. Given reward times *t*_*r*_ with probability distribution *p*(*t*_*r*_|*s*), the *η*-expectile 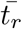 satisfies^71^:

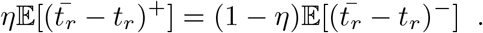

For example, the mean corresponds to the expectile with level *η* = 0.5. The expectiles are distributed according to the cumulative distribution,

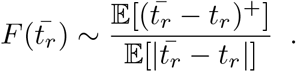

An explicit representation of this cumulative distribution has also been derived previously^72^.

Given a population activity vector of *N* dopamine neurons 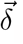, we sought to maximize the mutual information 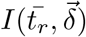 between the true expectile reward times 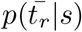 and those encoded by the population 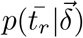, constraining on the number of neurons and on the population expected firing rate *R*. We consider a population with homogeneous derivatives,

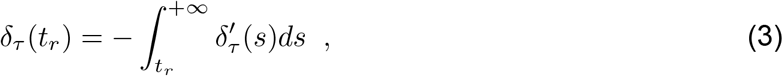

that approximately tiles the range of possible reward times from 0 to *T*, have constant Fisher information and where each neuron is subject to independent Poisson noise. A population that linearly tiles the exponential decay half-lives approximately satisfies these conditions, however this construction has edge effects at 0 which does not affect the capability of this model to encode future (non-zero) reward times. Since computing the mutual information is analytically intractable, a lower bound is optimized instead, namely the Fisher information^73^. We parameterize the optimized dopamine tuning curves _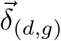_ with an invertible *density* function *d* : 𝒯 → 𝒯 which characterizes the heterogeneous allocation of neurons to reward time scales *τ* ∈ 𝒯 ≡ ℝ and a *gain* function *g* : *T* → *T* which characterizes the mean firing rate across reward times *t* ∈ 𝒯 ≡ ℝ,

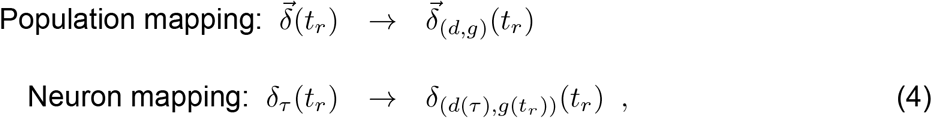

where *d*(*τ*) is the density of tuning curves at reward time scale *τ* in the optimized population and *g*(*t*_*r*_) is the population mean firing rate for reward time *t*_*r*_. The Fisher information of the optimized population is given by^22^,

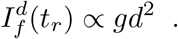

After constraining on the number of neurons *N*, the solution for the density function *d* is given by,

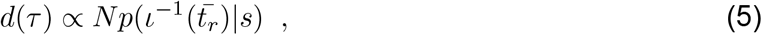

i.e., the population reward time scales should distribute proportionally to the expectile rewards times in the environment (see Ext. Data Fig. 10). On the other hand, additionally constraining on the mean population firing rate *R*, considering the population defined in Eqn. 3, the population gain function *g* should satisfy,

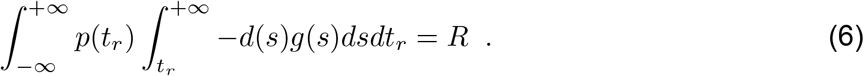

Integrating by parts to obtain the mean population firing rate as,

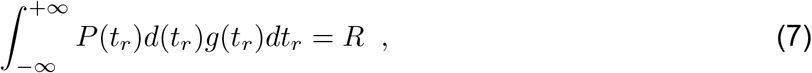

where *P* (*t*_*r*_) is the cumulative distribution of *p*(*t*_*r*_). Therefore the solution for the population gain is given by,

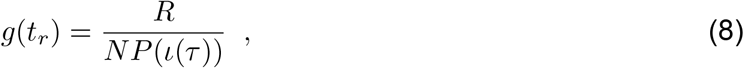

i.e., for each neuron the gain should be inversely proportional to the probability that a randomly chosen reward time will be smaller than it’s time scale. For small reward times, the entire population is active, incurring a large metabolic cost for encoding these values. Intuitively, this metabolic penalty can be reduced by lowering the gains tuned to long reward times and therefore large cumulative probabilities *P* (*ι*_*t*_(*τ*)) in Eqn. 8 (see Ext. Data Fig 10).

### 7.2 Distributional learning mechanism

In our analysis thus far, we have identified a particular density profile of neural tuning functions that optimally encode the distribution of reward times following a stimulus presentation. In this section, we describe how such a neural population may, in principle, be optimized via an online learning process.

We pursue an algorithmic strategy inspired by the distributional learning of reward magnitudes as proposed previously^7^ In this approach, multiple channels with different relative scaling for over- or under-estimation of reward magnitude (*α*^+^, *α*^−^) leads to a diversity of learned values,

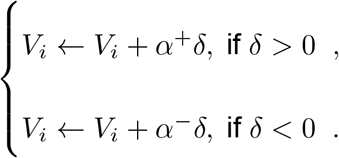

which collectively encode the entire reward magnitude distribution.

With respect to the distribution of reward times *p*(*t*_*r*_|*s*), we consider multiple channels with different relative scaling for over- and under-estimation of reward times thus generating a heterogeneous set of time scales. If *t*_*r*_ is the time of reward on a given trial, the corresponding prediction error is given by *δ* = *t*_*r*_ − *τ* and the update rules are

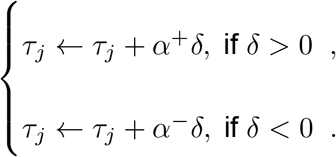

These learning rules converge to the *quantiles* of the probability distribution over expectile reward times 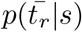^6^. This is because these quantiles are uniformly distributed in the cumulative probability space (i.e. the domain of the cumulative distribution function 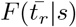 over expectile reward times, and satisfy the optimal information-theoretic condition defined above (Eqn. 5).

#### 7.3 Multi-dimensional integration over reward magnitude and time

Importantly, while in the distributional code for magnitude, the slope asymmetry 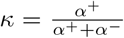 con-trols the level of *optimism* in individual units ^7^, in the temporal coding model developed here it controls the reward time scale and therefore the temporal discount factor, also known as *impatience*^74^. Integrating the distributional models in reward magnitude and time, leading to the time magnitude reinforcement learning (TMRL), each neuron is characterized by an optimism level *κ* and an impatience level *η*, corresponding to the temporal discount factor *γ*_*j*_ (which is induced from a corresponding reward time scale *τ*_*j*_). Therefore the temporal-difference vector RPEs take the form,

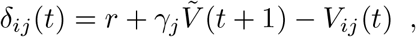

where 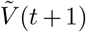 is a random sample from the value distribution at time *t* + 1. This is denoted by the *imputation* step, that implies a non-local update rule, as shown in previous work^70,75^. Understanding if, in the population of dopamine neurons, this manifests as a non-local update rule or may be implemented locally remains an open question. Finally, the multi-dimensional distributional value update rule is given by,

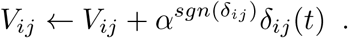

### 8 Data analysis

Spike counts were binned in 2-ms windows and smoothed by convolving with a gamma probability distribution kernel (shape parameter *k* =2 and scale parameter *θ* =25ms).

We identified putative dopamine neurons based on their firing rate patterns in a window around the cue and reward using an unsupervised clustering approach as in previous studies^76–18^. In summary, for each neuron and each 200ms time bin, the area under the receiver operator curve (auROC) was calculated between the distribution of firing rates across trials for that bin and the distribution of baseline firing rates (1s before cue). PCA of the auROC was calculated and then hierarchical clustering was done using the first three PCs using the Euclidean distance metric and the complete agglomeration method. As described before, three clusters were found: one revealed sustained inhibition at rewards (Type I), one had phasic responses to cue and reward, where the vast majority of photo-identified neurons were included (Type II) and the last had sustained excitation to reward (Type III). Type II neurons were classified as putative dopamine neurons. We focus our analysis on both light-identified dopamine neurons (n=43) and putative dopamine neurons (n=131).

Responses to reward were defined as the average activity from 200 to 650 ms after cue and reward onset, baseline subtracted by the mean activity over trials from -1000 ms to 0 ms relative to cue onset. Responses to cue were defined as the average activity from 200 to 650 ms after cue onset. This window was selected in order to exclude the initial response to the solenoid valve opening, that was in the majority of the neurons not selective to reward amount or delay, as shown in previous literature^79^. For each dopamine neuron *j*, we assumed the tuning function of reward time (*t*_*r*_) was given by an exponential decaying function^80^,

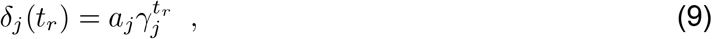

where *γ*_*j*_ is the temporal discount factor and *a*_*j*_ is a gain parameter. The parameters were estimated minimizing the mean squared error between *δ*_*j*_(*t*_*r*_) and the mean responses at the cue for each reward time.

To test whether the estimated temporal discount factors did not reflect noise we divided the trials in two random partitions, such that each partition contained the same number of trials for each reward delay. Then we estimated the temporal discounts for each partition and measured the linear regression slope, correlation coefficient and p-value for these sets of estimates. We repeated 10,000 times this procedure, and computed the mean correlation coefficient and the geometric mean of the obtained p-values.

In order to test if the single neuron’s estimated temporal discounts are significantly different from the population mean temporal discount, we took randomly selected 50% of trials per delay, estimated the temporal discounts and repeated 1000 times, to obtain confidence intervals.

To test if changes in the reward time statistics would lead to an adaptation of the temporal discounts, we removed either the longest or shortest delay and estimated the temporal discount before and after this context switch. The responses to the same delays were used to estimate the temporal discounts before and after the manipulation. After the context switch, we did not consider the first 5 trials. To assess the degree of relative adaptation of the population activity when removing the shortest or longest delay at the end of the session, we bootstrapped considering 10,000 resamples and computed the p-value and confidence interval (for the null hypothesis that there is no adaptation in the mean population temporal discounts). We also tested if there was adaptation of the population temporal discounts separately, for each type of context switch, bootstrapping using 10,000 resamples and computing the p-value and confidence interval (for the null hypothesis that there is no adaptation).

To test whether discounting by dopamine neurons was updated more continuously, we started by computing the rate of reward occurrence, by convolving the occurrence of rewards with exponential discounting kernels with different time scales. For different time scales we observed the lower the rates, the larger the population temporal discount factors. We represent the adaptation in temporal discounts for the time scale that minimized the bootstrapped p-value (considering 10,000 resamples) comparing the update in temporal discounts for low and high rates (Ext. Data Fig 9). Since for low reward rates the number of small reward delay trials was lower, and for high reward rates the number of high delay trials was lower, when fitting the temporal discounts factors, we weighted the errors by the normalized variance of the responses.

On the other hand, for each dopamine neuron *i*, we assumed the tuning function of reward magnitude (*r*) was given by a bilinear function,

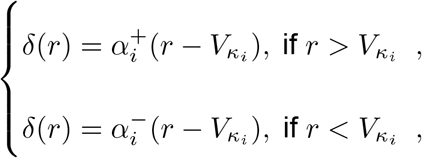

where 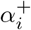 if the slope for the positive RPEs, 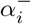 the slope for negative RPEs, 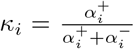 asymmetry in slopes for positive and negative RPEs and 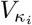is the reversal point. As in previous work^7^, the reversal point was defined as the magnitude *M* that maximized the number of positive responses to rewards greater than *M* plus the number of negative responses to rewards less than *M* . After measuring reversal points, we fit linear functions separately to the positive 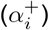 and negative 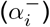 domains of each cell.

We measure the correlation between the firing rate at the cue corrected for the temporal discount and gain (Equation 9) and report the correlation coefficient, p-value and 95% confidence interval.

To test if the variance of estimated reversal points for the variable cues was significantly greater than for the certain one, we boostrapped considering 10,000 resamples and computed the p-value 95% confidence interval.

To classify an event as a lick it had to have a minimum duration of 0.015s. Licks were binned in 0.13s windows and smoothed by convolving with a exponential decaying function with a time scale of 0.77s to obtain lick rates. The anticipatory licking slope, described in Figure 3G, was defined as the linear regression slope in licking rate from *t* =0.01s to *t* = reward time.

### 9 Future reward distribution decoding

#### 9.1 Decoding future distribution of reward times

As defined before in Equation 10, the value considering a time horizon of T is given by,

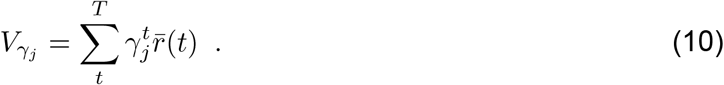

The problem of determining the expected future rewards over time 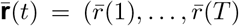 can be seen as a linear regression problem. The data points correspond to the population temporal discounts {*γ*_1_,. .., *γ*_*N*_ }, the basis functions are given by *φ*(*γ*)= (*γ, γ*^2^,…, *γ*^*T*^) and the targets are 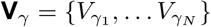, which are the single neuron’s responses at the cue. Assuming a uniform prior and a Gaussian likelihood functions with inverse variance *β*_*j*_ the log of the posterior is given by,

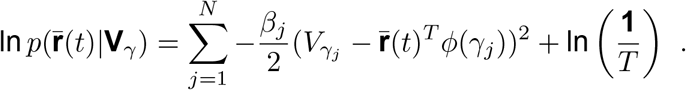

When the reward is either 0 or 1 the expected rewards at a given time-step *t* corresponds to the probability of rewards at that time-step, therefore the log posterior becomes,

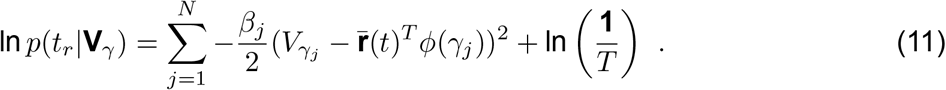

In practice, for each neuron we considered *V*_*j*_ the responses of each neuron normalized by the estimated gain and *β*_*j*_ was the estimated inverse of the variance of 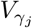 . The solution that maximized (11) was determined analytically using singular value decomposition (SVD), as proposed in^75,81^. To obtain a probability distribution, the solution was normalized. A qualitative search was done on the smoothing parameter, to maximize the similarity between the estimated and the true probability distribution.

For Figure 3E, we computed the mean decoded densities using the population responses in trials with anticipatory licking slope greater or smaller than the median.

#### 9.2 Decoding future distribution of rewards amounts

The population firing rate at the cue corrected for the gain and temporal discount factor 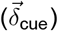 is significantly correlated with the reversal points estimated at the reward delivery. We therefore assume an additive Gaussian noise model,

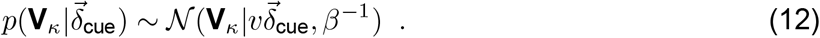

In practice, for each neuron, *β*_*i*_ was the inverse of the estimated variance of 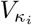 . Each dopamine neuron *i* has a different level of optimism *κ*_*i*_, by weighting asymmetrically positive 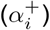 and negative 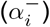 RPEs, with 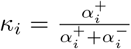, predicting different values, 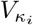. We measure the responses of each neuron at only five different reward amounts in a small range of rewards, from 1*μ*l-8*μ*l. We use Monte Carlo simulations to estimate the bias and variance we are inducing in the estimation of the *κ* parameter. For these simulations we assume the noise in the responses is Gaussian and use the variance of each neuron’s responses for each reward amount to generate responses. We observe that there is a sistematic relationship in the induced bias and variance: the variance increases quadratically with the distance to the mean of the reward magnitudes and the absolute bias increases linearly with the distance to the mean of the reward amounts, and it is positive or negative for reversal points smaller or greater than the mean reward amounts, respectively (Ext. Data Fig. 8). We therefore subtract the induced bias to the estimated *κ*_*i*_’s. We use a isotonic regression to model the relationship between the reversals *V*_*κ*_ ‘s and *κ*_*i*_’s^82^, which only assumes this is an increasing function. To estimate the piecewise linear functions in isotonic regression we take in account the variance in the estimation of each *κ*_*i*_. We assume the population of *N* neurons is minimizing the loss function^7^

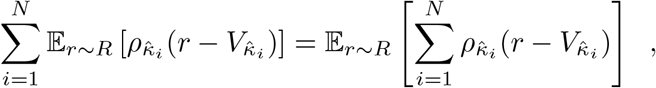

where

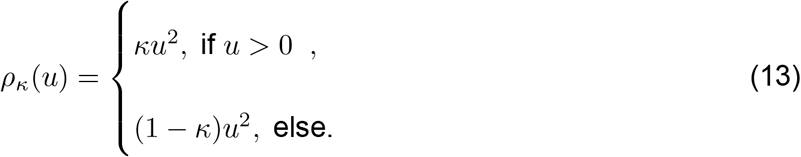

Considering asymmetric Gaussian likelihood functions^83^ with inverse variance *β*_*i*_, we obtain the posterior over reward amounts,

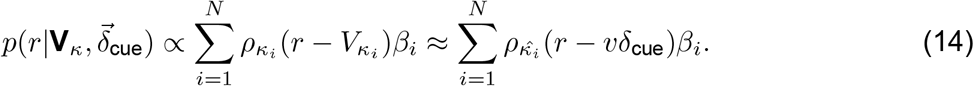

#### 9.3 Decoding future rewards over time and amount

Taking in account the dopamine population diversity in value prediction in terms of temporal discount **V**_*γ*_ and optimism level **V**_*κ*_, the posterior over reward magnitude and time is given by,

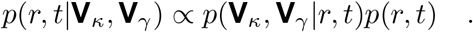

There is evidence that the diversity in dopamine tuning functions to reward time is independent of the diversity to reward magnitude (see Ext. Data Fig. 11A), hence we can factorize the above equation,

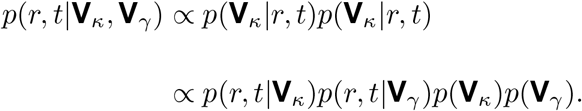

Conditioned on the cues, the amounts and times of reward are independent in our behavioral task, hence we assume the joint distribution over reward amounts and times can be factorized,

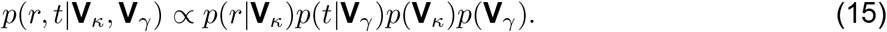

Thus we derive a probabilistic decoding framework for independently decoding reward time and magnitude. This, despite the limited numbers of neurons in our dataset, allows for the decoding of the reward distribution over these two dimensions in practice. In principle, with a sufficiently large number of neurons this assumption can be relaxed (see Ext. Data Fig. 11B,C).

### 10 Foraging simulations

In the foraging simulations in Figure 6 we implemented a temporal-difference learning algorithm for estimating the value of each patch (value RL). Simultaneously, we implemented a temporal-difference algorithm for estimating the occupancy matrix, that was multiplied by the reward, to obtain the sucessor representation (SR) estimation of value for each patch. For the TMRL we considered a set of values with different temporal discounts, and with these, we decoded the probability distribution over future reward for each patch. If the time of rewards in one patch was shorter, the agent selected that patch first. Initially the values for all algorithms were set to zero. The policy was the softmax over the value of each patch, with temperature parameter set to 0.2, however the simulations results are qualitatively independent of this parameter. A reference temporal discount was used to obtain a value and get a policy for TMRL. The learning rate was set to 0.02. For the stationary environment, patch one always retrieved a reward of 6 and patch two a reward of 2. For the non-stationary environment, patch one retrieved a reward of 2 after 2 time steps, patch two a reward of 0 or 4 after 2 time steps and patch three a reward of 6 after 6 time steps. Early in learning corresponds to 1000 time-steps and late in learning corresponds to 10,000 time-steps.

In the foraging simulations in Figure 6E, the change in internal state from hungry to sated was modelled as a change in the intrinsic value or utility of reward. We model the utility function in time as a decaying function in time parametrised by the temporal discount factor (*γ*^*t*^). When the mice was hungry the temporal discount was set to *γ* = 0.6 and the utility function in reward magnitude was considered convex^84^. When the mice became sated, the temporal discount factor was increased to *γ* = 1^85^ and the utility function in reward magnitudes was considered to be linear. We consider the utility matrix *U* defined over a discretized time range, {1,. .., *j*, . .., *T* }, and magnitude range, {*u*_1_,. .., *u*_*i*_,. .., *u*_*N*_ },

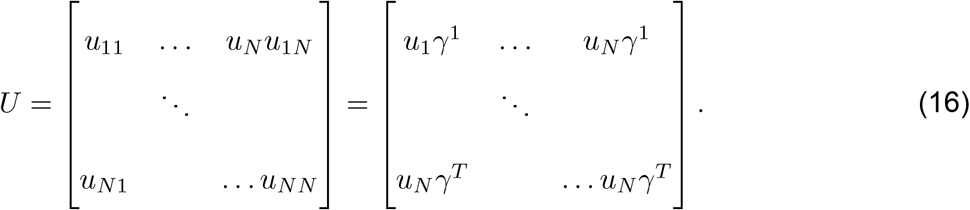

Since the TMRL allows for the construction of a probability distribution over future reward time and magnitude, once the agent has experienced reward in the three different patches for the sated state, it can recompute the value of each patch for the new state, using the distributional map. In particular, for each time-step in the future, the probability distribution over reward magnitudes *p*_*ij*_ is normalized, such that for all *j*,

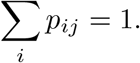

Then, each entry of this matrix is multiplied by the respective element in the utility matrix and summed to obtain the value for each patch,

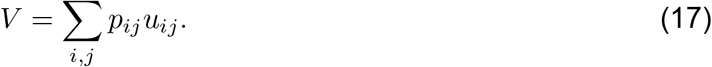

The SR agent can also flexibly recompute the value function for the sated state using the occupancy matrix. However, the standard value RL agent needs to learn through experience to compute the value of each patch for the new state.

## Extended data figures and tables

**Extended Data Figure 1:**
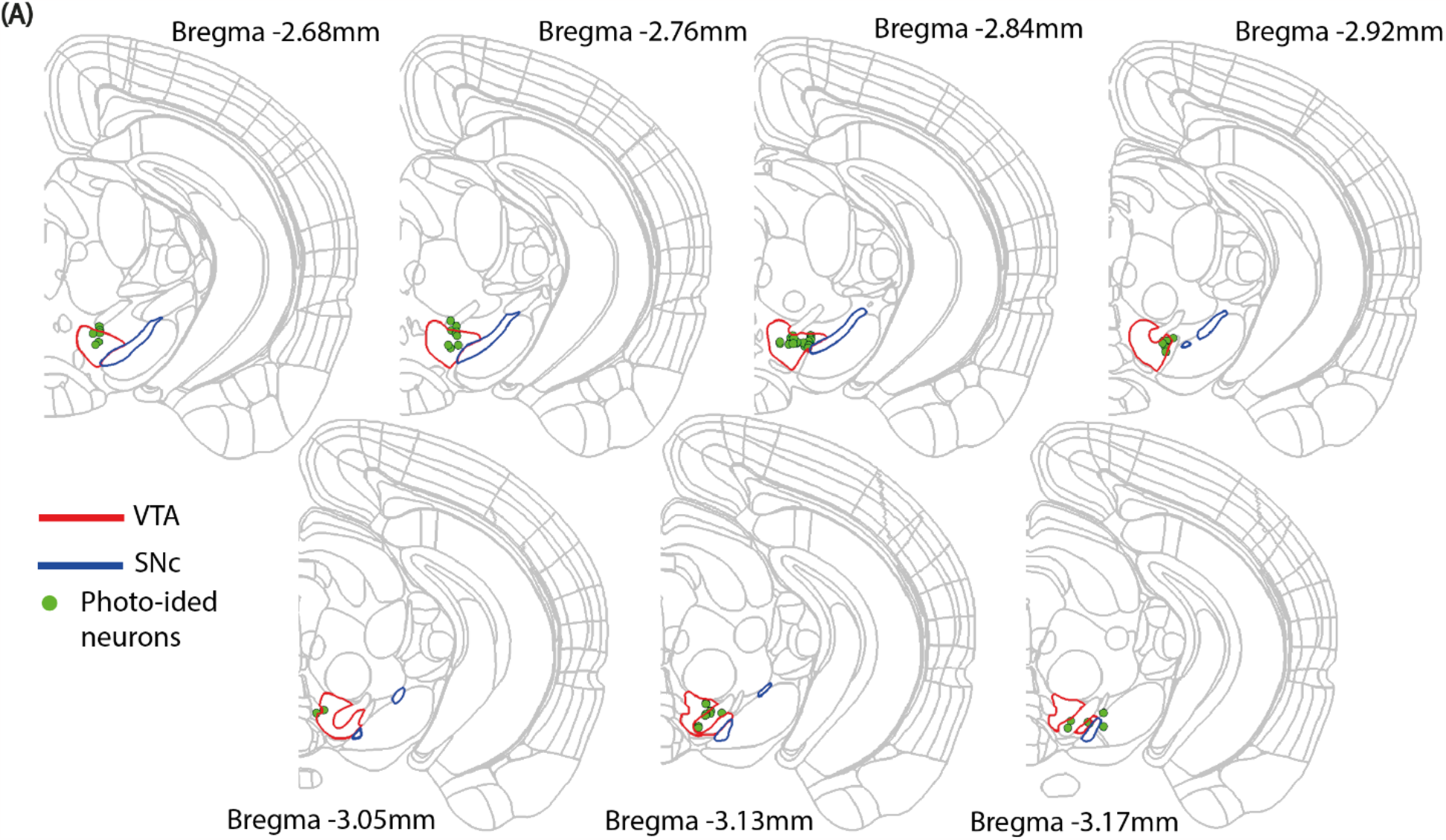
Histological reconstruction of recording sites. Recording locations of the probe tracks. Recording sites from all mice in seven coronal sections from the rostrocaudal axis (AP -2.68, -2.76, -2.84, -2.92, -3.05, -3.13,,−3.17). The approximate location of the photo-ided neurons is depicted in green, the VTA has a red outline and the SNc a blue outline.

**Extended Data Figure 2:**
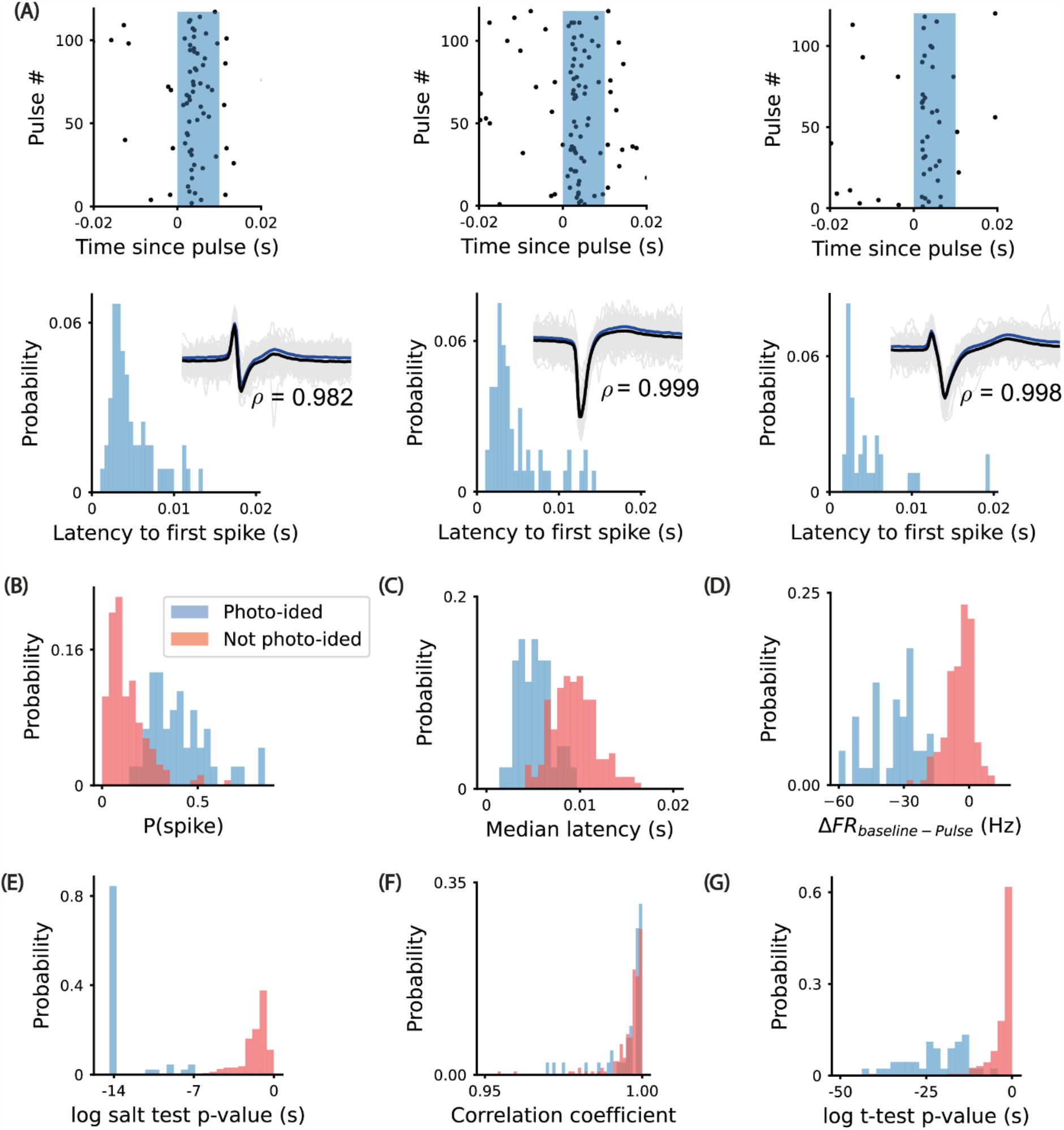
Photo-identification of dopamine neurons. **(A)** Three example photo-ided neurons. **Top)** Raster plot with single spikes aligned to laser pulse onset (10 ms duration). **Bottom)** Distribution of latencies to first spike after laser pulse observed in a 1-20ms window Bottom-inset) mean waveform (black) and mean laser-triggered waveform (blue). Distribution of: **(B)** probability of observing a spike xsbetween 1 and 10 ms after laser onset pulse. **(C)** median latency to first spike in a window between 1ms and 20ms. **(D)** Differences in firing rate between the baseline and 1-10ms post-pulse window. **(E)** Log of the p-value of the salt test^54^. **(F)** Correlation coefficient (*ρ*) between the mean waveform and the mean laser-triggered waveform. **(G)** Log of t-test for the difference in firing rates between baseline and pos-pulse firing rate in a window 1-10ms.

**Extended Data Figure 3:**
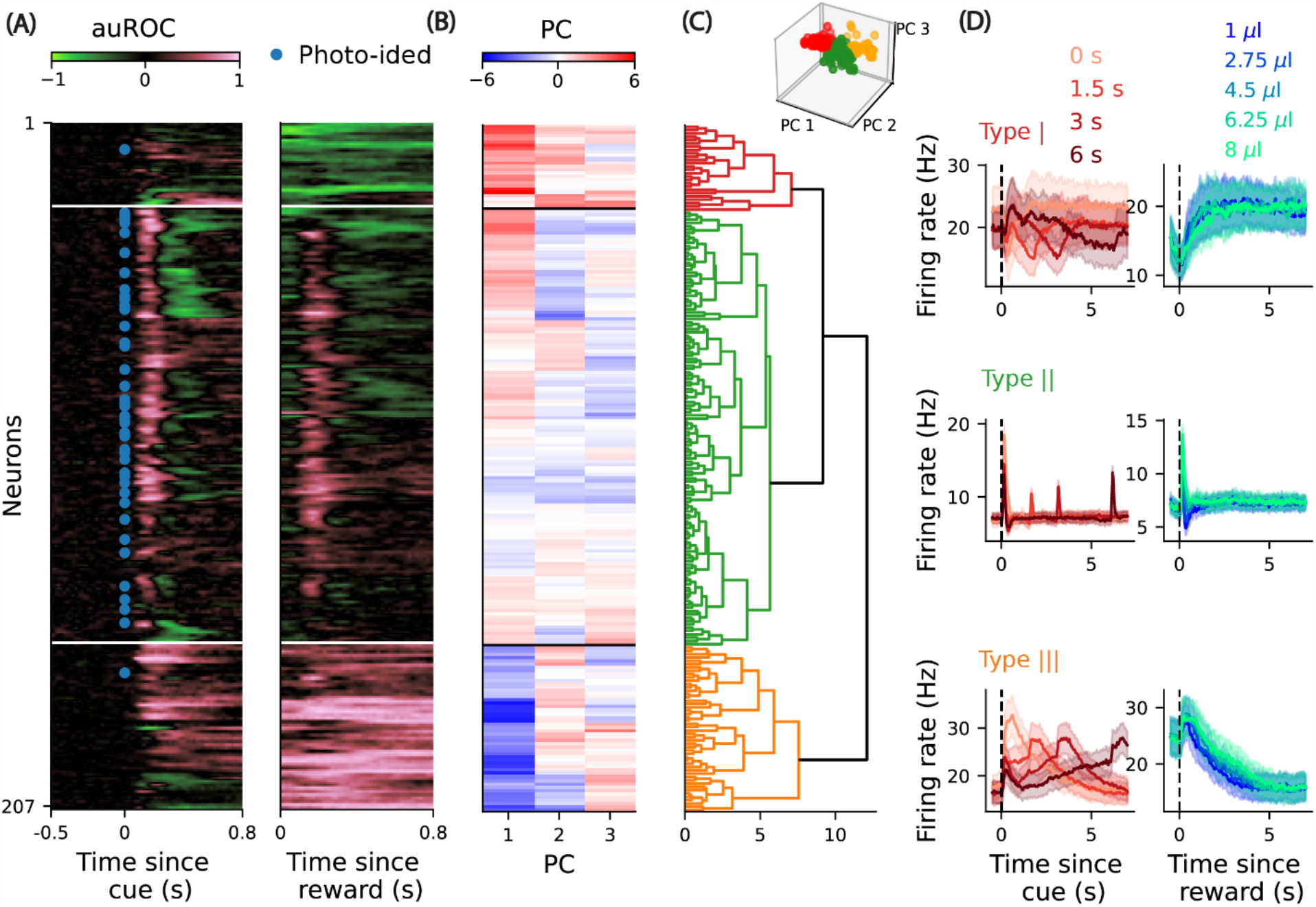
Hierarchical clustering of VTA/SNc neurons. **(A)** Area under the receiver operator curve (auROC) aligned to cue and reward delivery for the recorded population of neurons. Pink corresponds to increase relative to baseline activity, green to decrease and black to no change. The blue dots correspond to photo-ided neurons. **(B)** Projection coefficient for the three first principal components (PCs) of the auRoc. **(C)** Using these projection coefficients, hierarchical clustering with the euclidean distance was performed using the complete agglomeration method. **Top:** Each dot corresponds to a recorded neuron in the 3 first PC space. **(D) Left:** PSTH for each cluster for the different reward delays aligned to cue. **Right:** PSTH for each cluster for the different reward amounts aligned to reward delivery.

**Extended Data Figure 4:**
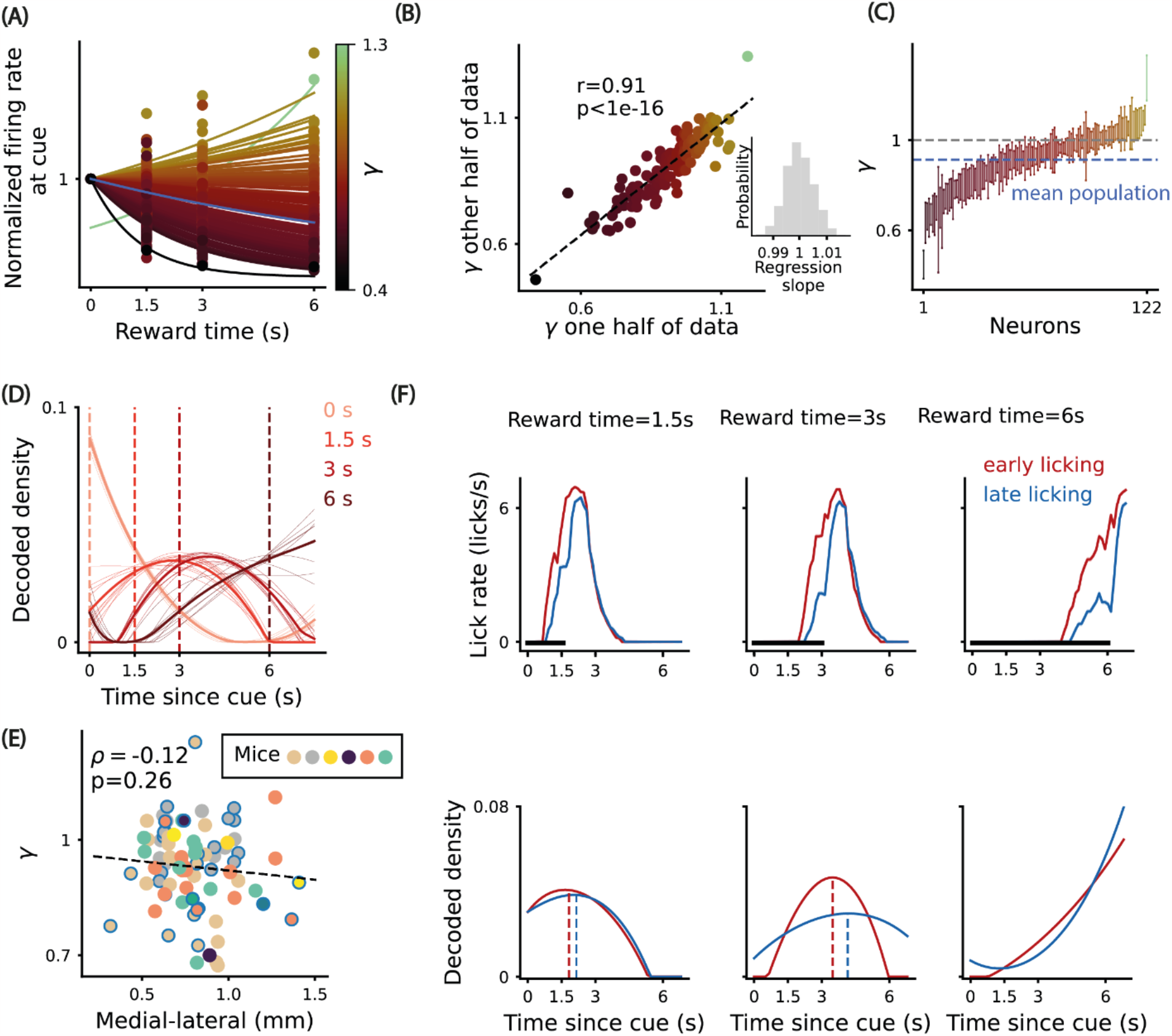
Heterogeneity in temporal discounting in putative dopamine neurons. **(A**,**B**,**C**,**D**,**F)** Analysis from Figure 3 with putative dopamine neurons. **(E)** Estimated single neuron temporal discount factors as a function of the position in the medial-lateral axis. Dots are color coded by animal identity and dots with a blue outer circle correspond to the photo-identified neurons. The line represents the Huber Regression fit. Spearman rank correlation, two-tailed test p-value=0.26.

**Extended Data Figure 5:**
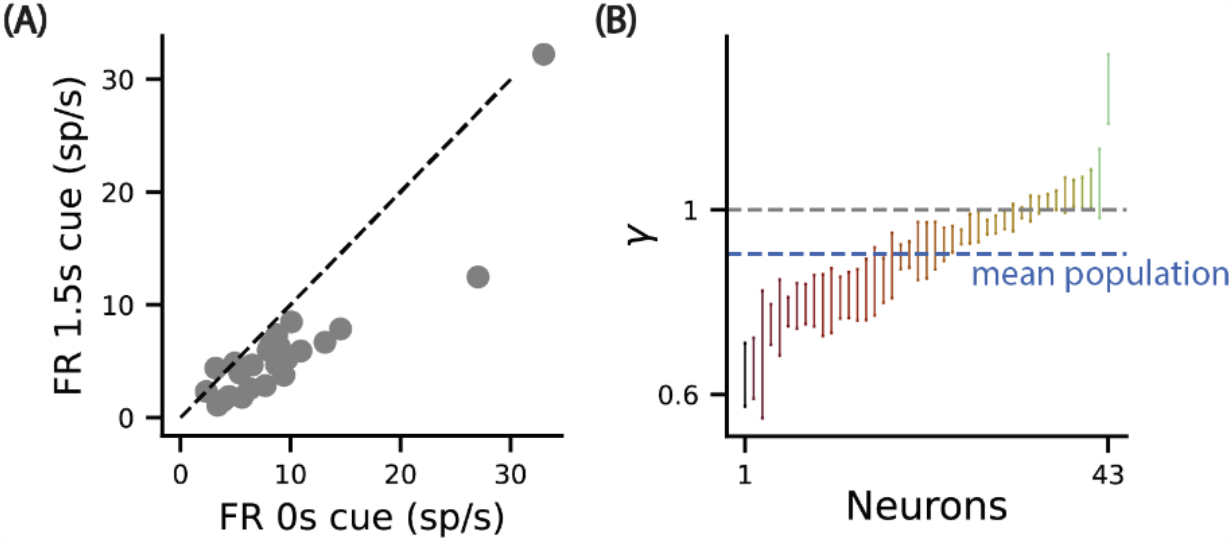
Testing the influence of number of experienced delays on estimated temporal discounts. **(A)** Responses at the 1.5s delay cue as a function of the responses at the 0s cue. The dashed line represents the unitary line. **(B)** Similar to Figure 3 (C) but considering the responses at the 1.5s reward as an (under) estimator of the 0s cue responses, for neurons recorded from the mice only subject to three delays.

**Extended Data Figure 6:**
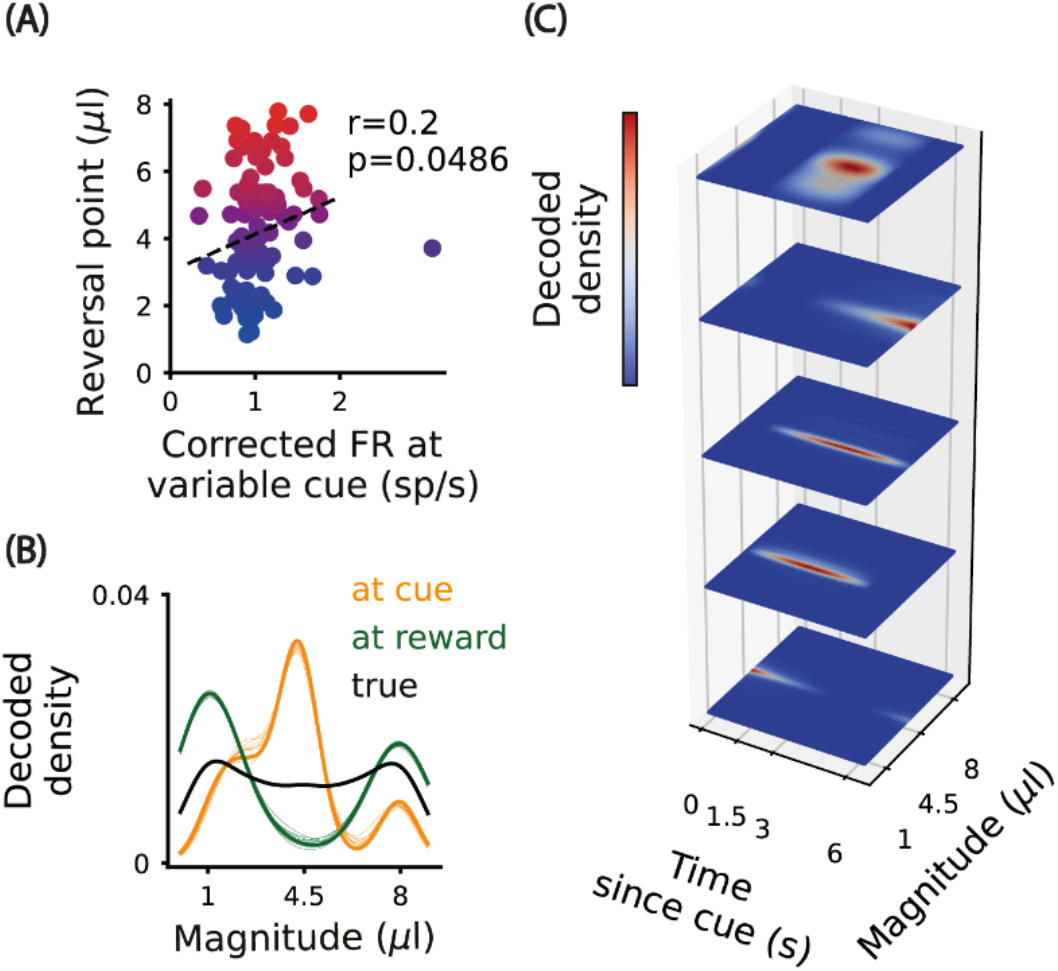
Analysis from Figure 4 with putative dopamine neurons.

**Extended Data Figure 7:**
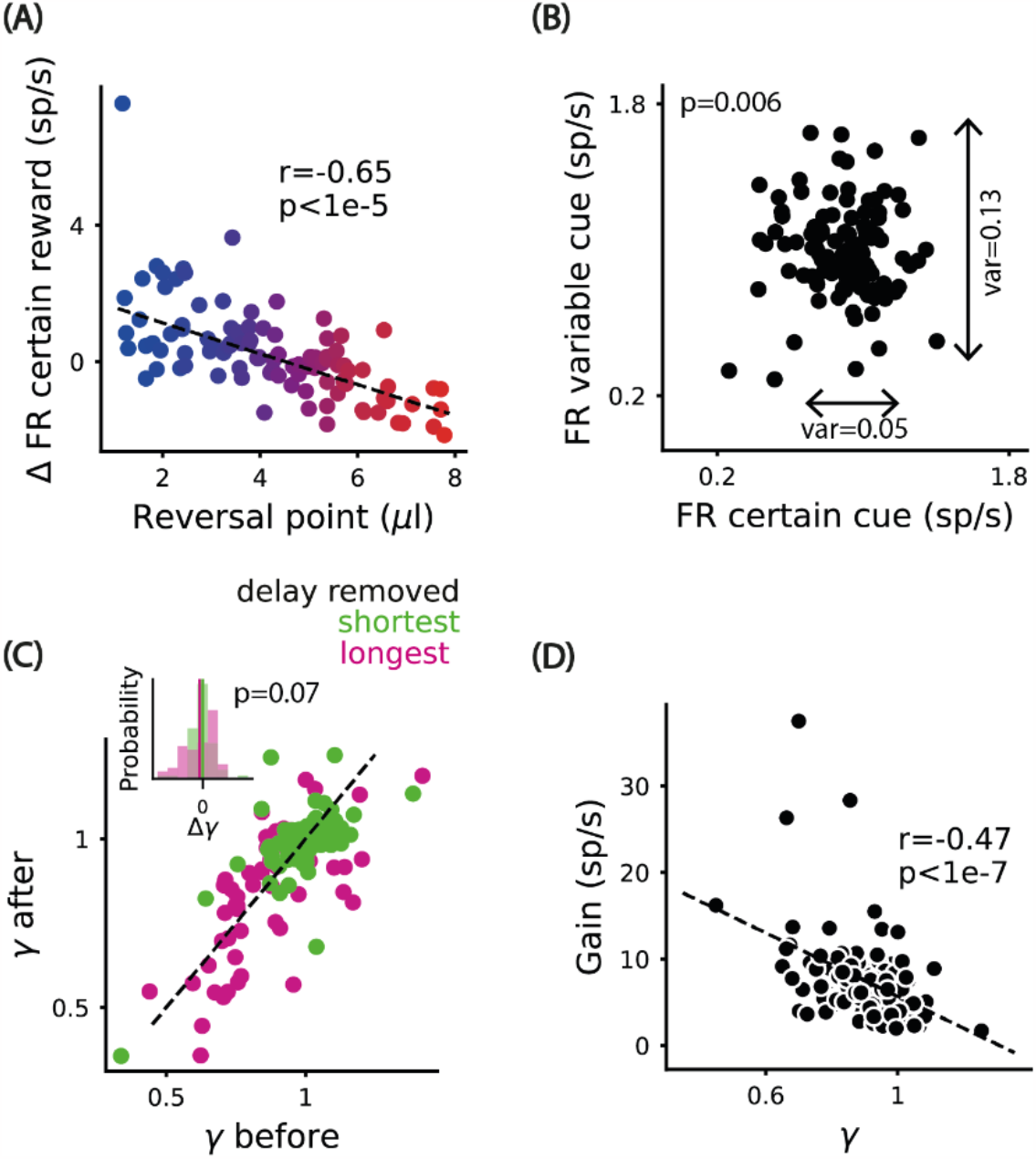
Analysis from Figure 5 with putative dopamine neurons.

**Extended Data Figure 8:**
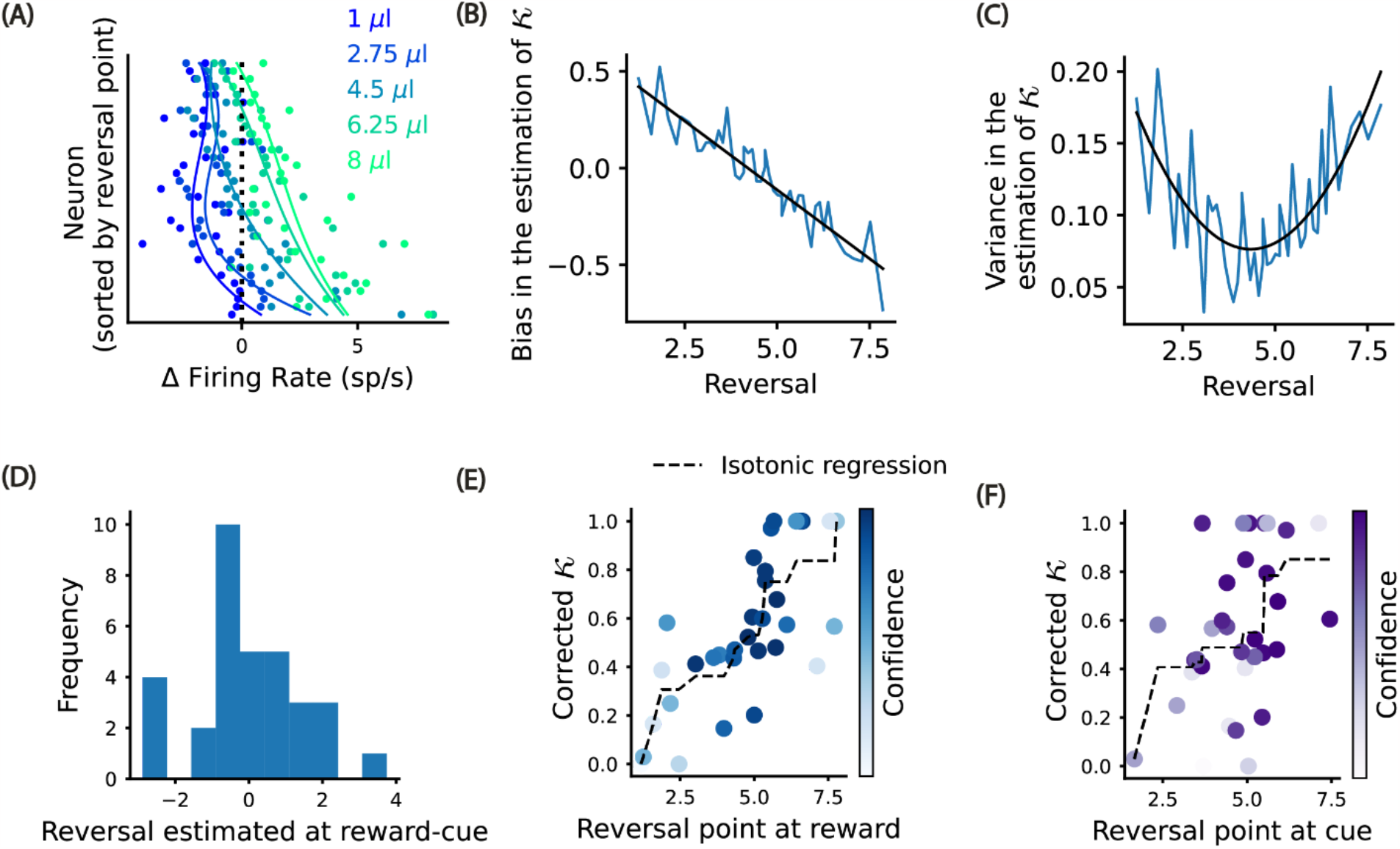
Decoding distribution over reward magnitudes. **(A)** Responses for different reward amounts for individual dopamine neurons sorted by estimated reversal point. **(B)** Bias in the estimation of as a function of reversal point, computed with monte carlo simulations. **(C)** Variance in the estimation of as a function of reversal point, computed with monte carlo simulations. **(D)** Distribution of the difference between the reversal point estimated at reward and at the cue, using the linear regression depicted in Figure 4A. **(E)** Corrected asymmetry as a function of the reversal point estimated at the reward, color coded by the confidence. The dashed line represents the isotonic regression fit. **(F)** Corrected asymmetry as a function of the reversal point estimated at the cue, color coded by the confidence. The dashed line corresponds to the isotonic regression fit.

**Extended Data Figure 9:**
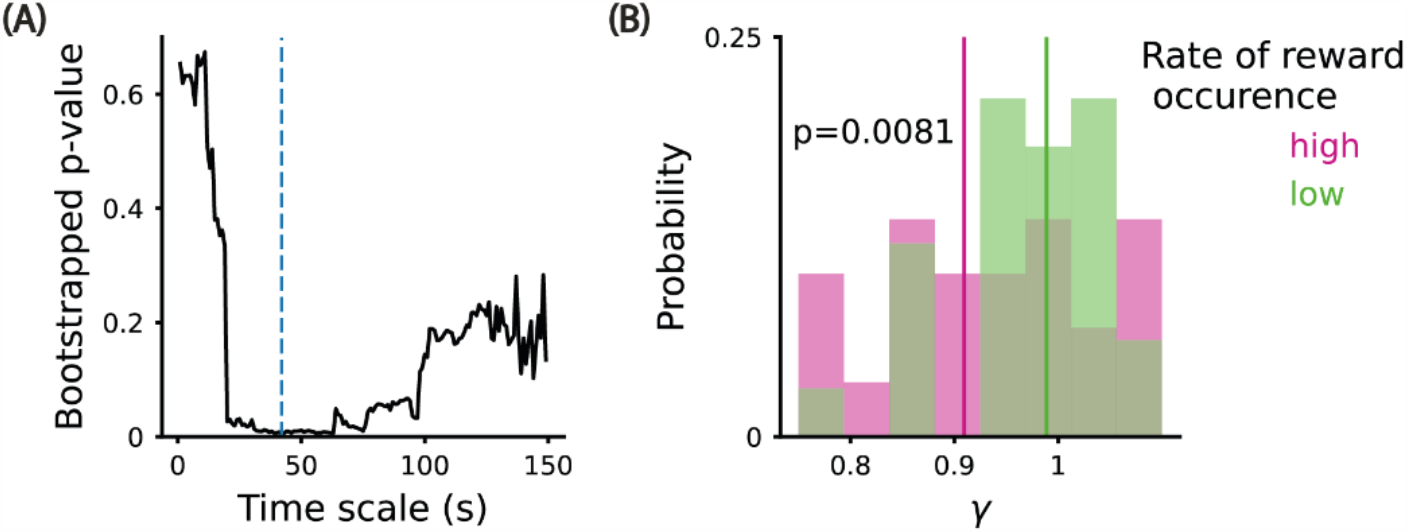
Adaptation of dopamine neuron population temporal discount factors with reward occurrence rate. **(A)** Considering exponential decaying kernels with different time constants to compute the rate of reward occurrence, we compute the p-value bootstrapping 10000 times, comparing the update in the estimated temporal discount factor for relatively high or low rate of reward occurrence. Vertical blue line: time scale of 39s that minimizes the p-value. **(B)** The estimated temporal discounts updates for relatively low or high rate of reward occurrence, computed using a exponential decaying kernel with the time scale that minimizes the p-value in (A). The correspondent 95% CI=(0.02,0.24). The vertical lines are the mean update in temporal discount rates.

**Extended Data Figure 10:**
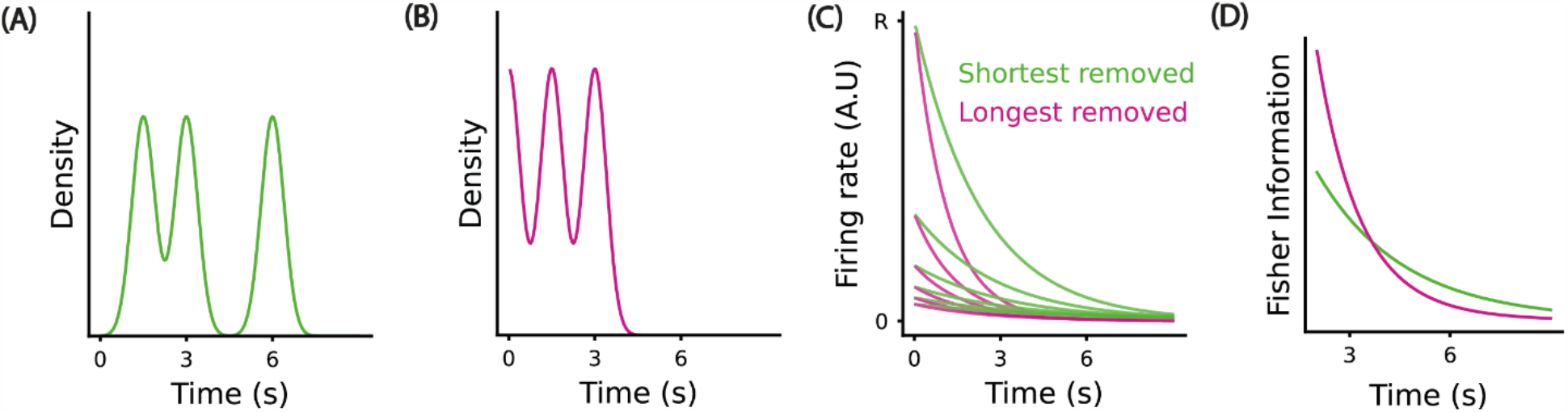
Efficient coding predictions. **(A**,**B)** Densities after removing the shortest (green) or longest (pink) reward delay. **(C)** The optimized tuning functions for the densities of reward time depicted in A and B. **(D)** The Fisher Information for the optimized populations for the range of 2-8 seconds.

**Extended Data Figure 11:**
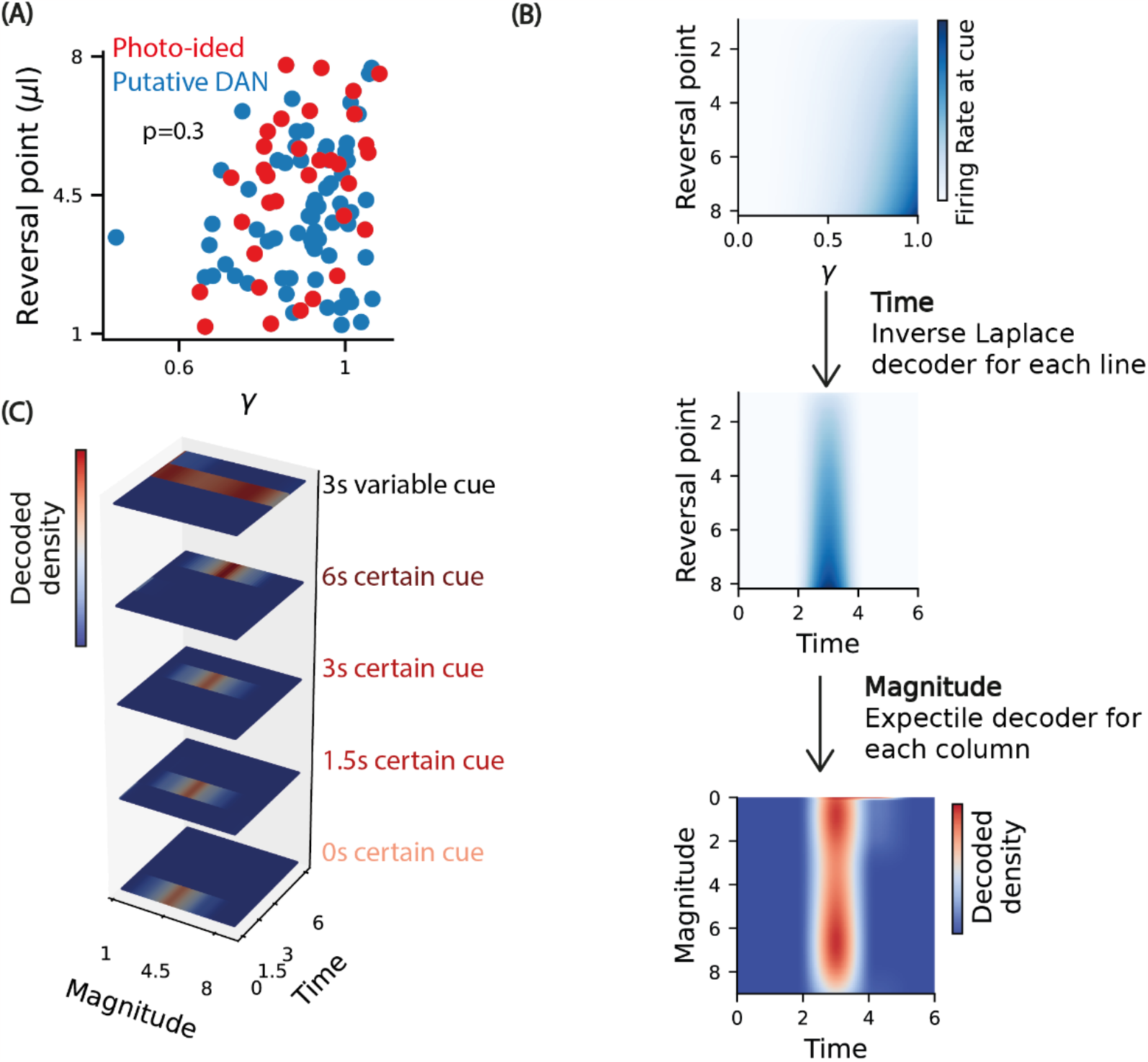
At a population level, the distribution over temporal discounts and reversal points are statistically independent. However, in principle, the joint distribution can be decoded without assuming factorization over time and magnitude. **(A)** Estimated reversal points as a function of estimated temporal discount factors, for photo-identified and putative dopamine neurons. Only neurons with reversal points in the range of reward magnitudes given in the experiment (1 μl-8μl) were included. The p-value refers to the chi-squared test, considering the null hypothesis that the joint distribution over the temporal discounts and reversal points is equal to the product of the marginals, and considering 16 degrees of freedom. **(B)** We simulate a population of n=100 units with a diverse set of temporal discount (γ) and reversal points uniformly sampled and decode the reward distribution over magnitude and time (not assuming these features are independent), from the responses at the cue. Importantly, these simulations were done for the cue that predicts a variable amount after a delay of three seconds, as described in Figure 2. We first apply the inverse Laplace for each reversal point and get the temporal evolution of reversal points (middle). Then, for each time we decode the distribution over reward magnitudes (lower), by scaling the probability for each time point by one minus of probability of reward=0. **(C)** The same simulations but for all cues described in Figure 4.

